# FPmotion an Automated Signal Processing and Statistical Analysis Tool for Fiber Photometry Data

**DOI:** 10.64898/2026.01.19.700261

**Authors:** Y. Hong, J.M. John, Thomas Topilko, P. Hollos, E. Coffey

**Author notes:** Corresponding author: Eleanor Coffey Ye Hong. FPmotion and FPviewer.exe softwares can be downloaded here: https://github.com/TCB-yehong/FPmotion.

## Abstract

Fiber photometry measures neural activity *in vivo* from genetically encoded indicators. Recordings generate large datasets that require extensive preprocessing and robust statistical methods for meaningful interpretation. However, existing analysis tools demand programming expertise, limiting accessibility. Here we describe *FPmotion*, a comprehensive, user-friendly software platform for batchwise processing, integration, and statistical analysis of fiber photometry data with or without accompanying behavioral information. FPmotion performs filtering, isosbestic regression, ΔF/F computation, detrending, and z-scoring. It also extracts detailed peak properties at whole-file, block-level, and individual-peak resolution. When behavioral data is provided, FPmotion automatically identifies behavioral bouts, computes bout-level statistics, performs peri-event signal extraction, and supports multi-group comparisons through ANOVA- and LMEM-based statistical frameworks. All analyses generate publication-ready figures and structured CSV outputs suitable for downstream workflows. FPmotion also introduces a dedicated alignment module for peri-event signal analysis. This module applies dynamic time warping followed by barycenter averaging to realign peri-event traces while preserving the behavioral time anchor, producing cleaner and more temporally coherent peri-event motifs. We demonstrate FPmotion’s capabilities using the dlight1.1 sensor to measure dopamine responses from ventral striatum in behaving mice before and after amphetamine treatment. Together, FPmotion offers a fully automated framework for FP data analysis that improves interpretability and accessibility while reducing analysis time substantially.

## Introduction

Fiber photometry (FP) enables real-time measurement of fluorescence reporter activity in specific brain regions of awake behaving mice (1). Initially used to monitor calcium transients from genetically encoded GCaMP reporters, FP facilitates investigation of neuronal circuit dynamics (1, 2). Chronically implanted optical fibers (200-400 µm in diameter) deliver excitation light to defined brain coordinates, enabling 1000 Hz acquisition with around 200 µm axial sensitivity, dependent on reporter expression and brightness (1, 3). A variety of fluorescent reporters are now used including dLight for dopamine, GRABNA for norepinephrine, iGluSnFR for glutamate, GABASnFR for GABA, in addition to serotonin, endocannabinoid and voltage sensors (4–7). The method is increasingly used to understand how neurotransmission and signalling in the nervous system control behavioural responses. Existing tools for FP analysis GuPPy (8), pMAT (9), pyPhotometry (10), FiPhA (11), Pyfiber (12), and PhAT (13), can be used to process the data and decode the underlying signals. However, these software packages do not provide built-in, robust statistical analysis combined with batch processing. Moreover, coding knowledge is required to run these pipelines. For this reason, FP has remained somewhat a niche method and the data analysis remains time consuming.

FPmotion is an FP analysis software package with G.U.I. that offers flexibility to the user. It is packaged entirely within an .exe file that can be downloaded and executed directly in a Windows environment. The user can choose to compare groups of differently treated mice during the analysis setup, so that in the output, the statistically significant changes are already annotated and indicated, both in csv and graphical formats, across a range of measurements and output files. Options for signal normalisation setup, applied either across animals or individually are available. This supports identification of patterns that are consistent across subjects, or accurate detection of stimulus responses within an individual. In addition to these features, FPmotion introduces a dedicated alignment module for peri-event signal analysis. This method uses dynamic time warping (DTW) combined with barycenter averaging to correct latency jitter across trials while preserving the behavioral time anchor, enabling cleaner visualization and quantification of peri-event motifs. To our knowledge, barycentering-based alignment is not included in existing FP workflows. Together with these analytical capabilities, FPmotion also emphasizes accessibility. The user-friendly GUI built with ttkbootstrap simplifies the analysis process, eliminating the need for coding and making sophisticated data analysis accessible to a broader range of researchers.

## 2. Materials and Methods

### 2.1 Introduction to the Software

FPmotion is distributed as a standalone Windows executable that can be downloaded directly from GitHub (https://github.com/TCB-yehong/FPmotion) and run without additional installations or programming expertise. The software provides an accessible, clearly structured graphical interface that guides users through the analysis in an intuitive step-by-step format, enabling users to begin analysis immediately after download. The software interface comprises four sequential sections that unlock progressively as each previous section is set up. The input interface accepts signal files and optional behavior and video files in bulk. The parameter interface provides customization settings for the analysis and output. The information input interface collects file information necessary for statistical methods and settings not covered in the parameter interface. The group comparison interface allows users to set up multiple groups and comparisons from the input files. Detailed descriptions and demonstrations are available in the accompanying vignette.

#### File naming

To match signal files with their respective behavior files and extract information directly from file names, we implemented specific naming conventions. The signal file name consists of six information fragments connected by underscores or spaces: “Trial_X” (trial number), mouse ID, genotype, behavior test name, and two experiment conditions. For example, “Trial_1_1827_control_OF_before_MK801” indicates treatment (“MK801”) and time segment condition (“before”). Behavior and video files only need to match the trial number with the signal files. Manual adjustments are possible if naming conventions are not followed.

#### Optional video file

Users can include mouse behavior videos in the analysis with any of the following format: mp4, .mov, .avi, .webm, .ogv, .mkv, or .gif,. If selected, additional video outputs align mouse movement, fiber photometry signals, and behavior bouts in the video. This option is time-consuming and is opted out by default but can be enabled in the parameter interface.

#### Fiber photometry signal processing

The default source for signal input files is Doric Lenses (Québec, Canada), although output from Tucker-Davis Technologies (TDT) and RWD Life Science (RWD) devices can also be directly imported into FPmotion. For signals from other sources, the required format includes three columns: the first for time points, the second for the isosbestic channel, and the third for the dopamine/calcium channel, saved in a .csv file. Raw fiber photometry signals undergo initial filtering. If the “dopamine” signal is selected, a Butterworth lowpass filter is applied, with customizable cutoff frequency and filter order. For “calcium” signals, a zero-phase moving average filter is applied in both forward and backward directions, with a customizable filter window. These filters are independently applied to the isosbestic and dopamine/calcium channels.

The isosbestic channel is then fitted to the dopamine/calcium channel using least squares linear regression. The percent dF/F is calculated by subtracting the fitted isosbestic channel from the dopamine/calcium channel, dividing by the fitted isosbestic channel, and multiplying by 100%. This % dF/F signal is detrended by subtracting a linear least-squares fit. Users can choose to calculate either a standard Z-score or an “external” Z-score. The standard Z-score is computed by subtracting the signal mean and dividing by the standard deviation. For the external Z-score, users can specify a set of baseline files and/or time periods; the mean and standard deviation of the % dF/F from these baseline signals are used in the calculation. All these calculations are performed using Python packages such as SciPy, sklearn, and NumPy. In addition, the following Python modules are used: OS, pickle, traceback, datetime, types, tkinter, ttkbootstrap, pandas, re, math, colorsys, webcolors, threading, peakutils, moviepy, matplotlib, seaborn, pingouin, statistics, statannot, csv, textwrap, scipy, sklearn, itertools, statsmodels, functools, pygam, tslearn, fastdtw, joblib and multiprocessing.

#### Behavior information processing

The default source for behavior information is output from EthoVision XT tracking software. For other sources, the input file should be in .xlsx format and have a column named “Recording time” for the behavior test time points, and subsequent columns for each behavior, with values of 0 (behavior not present) or 1 (behavior present). In the parameter interface, the behavior starting row should be set to 1. Each behavior is processed into behavior bouts, defined as continuous periods of 1s. The start time, end time, and duration of each bout are recorded. Bouts are adjusted to remove artifacts from signal processing, and those closer than a user-defined merging distance are combined. Bouts shorter than a user-defined minimum duration are excluded. The behavior syllable labels are automatically detected by Fpmotion from within the uploaded .csv file during the initial processing (see Supplementary User Guide, page 6, information interface I, point 2). The user can select one or multiple behaviors to analyse.

#### Combining Signal and Behavior - Combining signal and behavior

For each processed behavior bout, the lengths and counts per recording session are calculated and visualized with histogram overlaying kernel density estimates. Signal z-scores before and after each bout’s start are extracted, with each time segment defaulting to 5 seconds (customizable by the user). Peak thresholds define signal peaks, with customizable relative and absolute thresholds. The relative threshold uses the median absolute deviation of the signal z-scores multiplied by 2.91, while the absolute threshold is user-defined. A default peak distance threshold, defined as 100 data points, can also be customized. Once peaks are identified, their amplitude and frequency are calculated. Bar plots compare AUC, amplitude, and frequency before and after behavior bouts for each input file.

Single peak properties, such as time point, amplitude, width, prominence, and AUC, are categorized as within behavior bouts, outside of behavior bouts, and all passing signal peaks. This data is accumulated for statistical analysis on bulk data. Signals are also divided into shorter blocks to capture momentary variations, with a default block length of one minute. Each block inherits behavior bout and peak property information within its timeframe. Block-level data, along with behavior bout and peak property information, are output in CSV format for each input file. Signal peaks are extracted using Python’s SciPy package.

#### Group comparison utilizing bulk data

After individual file processing, data from all files are combined and saved in CSV files. Users can customize groups and comparisons in the group comparison interface, specifying which files belong to each group and which groups to compare. For behavior bout length and count values, statistical comparisons across groups utilize ANOVA with False Discovery Rate (FDR) corrections are performed. The results are visualized with detailed boxplots. For before/after behavior bouts data, grouped bar plots are created for amplitude, frequency, and AUC, showing pre- and post-bout measurements by comparison groups. A 2-way ANOVA with post hoc Tukey test is conducted to assess the significance of comparison groups and behavior effects. Pooled behavior bout data are visualized in heatmaps. For each pre- and post-bout signal combination with user chosen lengths, an optional outlier removal step can be performed by detecting and removing signals that are statistical outliers using robust covariance methods. Subsequently, signals are grouped into clusters based on their shape and temporal pattern similarity using Dynamic Time Warping (DTW) (14, 15). Once clustered, signals within each group are precisely aligned around the behavior events using DTW-based averaging (barycenter alignment). Alignment is performed separately for periods before and after behavioral events if chosen. The aligned signals are then analyzed for their peak properties, such as amplitude and prominence, and visualized in bar plots and violin plots. Signal peak data are analyzed on three levels: whole file average, block average, and individual peak level. For whole file averages, each peak property’s average value per file is used in statistical tests and visualized in bar plots with T-test significance. For block averages, average values are calculated per block, and line plots display these averages by group. For individual peaks, signal peak data are analyzed using linear mixed-effects models (LMEM) to account for within-subject variability and repeated measures. The resulting fixed-effects summaries, predictions, and statistical output are recorded, and visualized in scatterplots.

#### Two conditions comparison

A comparison setup allows optional comparison of two conditions from the file information: genotype and one other condition (e.g., drug treatment). This feature can be activated through the GUI’s parameter interface under ’General’ settings when relevant. Users can switch the genotype information if needed.

Segment line plots and scatter plots are generated for time block and single peak data of peak properties, segmented by the first condition and grouped by genotype. ANOVA tests, including 3-way ANOVA and 2-way ANOVA with post hoc Tukey tests, assess the significance of genotype, behavior bout presence, and the first condition, along with their interactions. Figures and statistical tests are created using Python packages seaborn, statannot, analys, and pingouin.

#### Analysis without behavioral input

Although FPmotion is designed to integrate behavioral tracking with fiber photometry data, users may also perform the full signal-based analysis without supplying behavior files. In this mode, all behavior-dependent steps—including bout detection, bout length and count summaries, peri-event signal extraction, and behavior-segmented analyses at file, block, and single-peak levels—are omitted. All signal-centric analyses remain available. This includes peak preprocessing, peak detection and characterization, and all statistical comparisons involving peak amplitude, frequency, width, prominence, and AUC at whole-file level, block-partition level, and single-peak level. This option enables users working with simple photometry recordings, pilot datasets, or experiments without behavioral annotation to obtain comprehensive signal-based results.

#### Peak alignment and outlier removal

FPmotion includes a dedicated module to enhance peri-event fiber photometry signal interpretation, before and after a behavior. Although all events in our datasets are time-locked to the annotated behavioral timestamps, peaks and troughs that are typically temporally misaligned across trials, lead to flattening and masking of physiologically relevant transients. To this overcome limitation, we implemented a multi-step workflow consisting of robust outlier removal, time-series clustering, and dynamic time-warping (DTW)-based alignment to a consensus template (16). Outlier detection was performed using the Mahalanobis distance, which is well-suited for high-dimensional time-series data. However, directly applying Mahalanobis distance to raw signals proved unstable due to singular covariance matrices, as signals often contain thousands of timepoints but far fewer trials. To address this, each signal was projected onto a reduced principal component (PCA) space, retaining *N–1* principal components for *N* trials (17). This ensured that the dimensionality never exceeded the number of observations. To prevent minor principal components, which primarily capture noise, from disproportionately influencing the distance metric, each component is normalized by its explained variance, effectively weighting the features in proportion to their contribution to overall signal structure. A robust covariance estimate is then obtained using the Minimum Covariance Determinant (MCD) method, which is less sensitive to extreme signals (18, 19). Outliers were identified by comparing principal component-space distances to a chi-squared threshold determined by the number of retained components. This approach consistently removes anomalous signals that deviate strongly from the dominant temporal structure, enabling cleaner downstream modelling.

Even after outlier removal, the signal ensemble can contain multiple distinct temporal motifs depending on behavioral context or neural heterogeneity. To assess whether the dataset naturally partitions into discreet patterns, we incorporated an unsurpervised clustering step. Because clustering raw signals is confounded by onset jitter and latency differences, we first computed dynamic time warping (DTW) distances between all second-half segments (the behavior-evoked portion of the signal). DTW was chosen because Euclidean distance cannot account for temporal misalignment of peaks or valleys, which is especially relevant when physiological latencies differ across trials despite shared behavioral timing.

Classical DTW is computationally prohibitive for datasets of this size (20), so we used fast DTW, an approximate implementation that provides strong performance with substantially reduced runtime. Parallel processing further accelerates distance computations, with our software automatically detecting available CPU cores and allocating two-thirds of them to the DTW process. We also apply time-series k-means clustering using the precomputed DTW distance matrix and evaluate clustering performance using silhouette scores. Other indices, such as Davies–Bouldin and Calinski–Harabasz were tested but removed due to excessive computational cost under DTW metrics. To guard against spurious structure, we generate random cluster assignments and recompute silhouette scores as a null baseline. A cluster solution is accepted only if its silhouette score exceeds the random baseline by more than two standard deviations; otherwise, all trials are treated as a single group. In practice, clusters containing only a few signals typically represent residual noise and are considered outlier-dominated. This step ensures that alignment is not forced into artificial partitions and preserves the integrity of behavior-linked signal motifs.

#### FPviewer: a results browser for FPmotion output)

To facilitate navigation of the large number of output files generated during bulk processing, we developed a standalone results viewer for FPmotion. This .exe file can be downloaded from here: https://github.com/TCB-yehong/FPmotion. The FPviewer organizes outputs into structured tabs corresponding to major result types: file overview, results from group comparisons for behavior bouts, file-level signal summaries, block-level signal properties, peak-level statistics, and before/after behavior segment analyses. For proper visualization of this structure, users should select the main results directory rather than an individual subfolder.

FPviewer will automatically map the folder hierarchy into its tab-based layout. Both image and text-based results can be previewed directly within the viewer, allowing users to browse heatmaps, bar plots, violin plots, and CSV-based statistical tables without opening files individually. Selecting a file from the tab displays its content in the central viewing panel, and a dedicated button allows users to open the displayed file externally when preparing figures or extracting values for manuscripts and presentations.

### 2.2 Mouse surgeries and fiberphotometry

#### Mice

Adult (8–12-week-old male) C57BL/6NCrl wild-type mice were maintained on a 12 h light/dark cycle and supplied with food and water ad libitum. Behavioural tests were performed between 08:00 and 15:00 h. The number of animals used in each experiment is indicated in the legends. Ethical permission: The animal work was done according to the project permit (ESAVI/30686/2022 and ESAVI/26777/2023) granted by the Regional administrative Agency of Finland.

#### Stereotaxic surgery

Individually housed mice were weighed and anesthetized with isofluorane (Isoflurane concentration in breathing zone of 2.5% for induction and 2-1% maintenance) and placed on a stereotaxic frame (KOPF, US) for the surgery. The depth of anaesthesia was estimated using the paw or tail response. A heating pad (Physitemp MTC-1, temperature controller at 37 °C) was used during the surgery to keep mice warm. Local anaesthesia 1mg/kg Lidocaine (Orion Pharma) in 125µl was administered subcutaneously at the incision point 2 minutes prior to making the incision. Temgesic (Indivior) 0.1mg/kg (170µl for a 25 g mouse) was given intra-peritoneally just before the surgery. The head was straightened and fixed on the frame. Viscotears were applied to prevent eyes from drying. An incision was made along the midline, the scalp retracted, and the area surrounding bregma was cleaned and dried. Coordinates (+1.54 mm AP, +/-0.6 mm ML, −4.5 mm DV) were marked carefully on the head and holes are drilled using the dentist’s drill (FOREDOM, K.1070). Infusion with pAAV-CAG-dLight1.1 viruses (0.2µl, Plasmid #111067 from Addgene) was performed using an automated sterotaxic injector (#53311, Stoelting, IL, US) at a rate of 0.10 µl/min and the needle was left in place for 10 minutes to prevent back flow.

Fiber optic cannulas (Doric lenses, core diameter 200 µm, code-200/230-0.48, MF1.25-X mono fiber optic cannulas) are then placed in the exact coordinates using the cannula holder (Thorlabs, SE) and fixed using dental cement (Super-bond C&B, Sun medicals, Japan). The wounds are stitched and saline 10 ml/kg is injected intra-peritoneally. After surgery, the animals are transferred to their “heat-pad warmed” home-cage to recover. Mice are monitored twice daily during the recovery period. Following surgeries, mice are housed alone for at least 2 weeks before behavioural tests are conducted. This period allows ample expression of virus.

#### Fiber photometry

Fiber photometry data was collected using a Doric fiber photometry customised system with optical components from Doric Lenses (Quebec, CA). For recording from ventral striatum using dLight1.1. FP measurements were acquired with an analog/digital acquisition console using LED excitation for dLight1.1 through an LED driver with 488 nm and background 530 nm. Excitation power was set between 40-60 µW and measured from the flat-end optic fiber. Emission was recorded at 1000 Hz sampling rate using Newport 2151 Femtowatt photoreceivers (Irvine, CA, US) in ‘DC-LO’ mode. Before connecting the fibers to the mice, fibers were photo-bleached for 10 to 15 minutes using 500 mA current in both the LED channels. After head mounting the fiber, *in vivo* autofluorescence from tissue was bleached for 5 minutes with 60µW before the experiments commenced. Behaviour tracking and FP acquisition were synchronised using a TTL trigger from Noldus I/O box (The Netherlands) connected to the FP console. Raw signals were recorded with Doric Neuroscience studio software and thereafter analysis was done using FPmotion software. The following settings were used for the FPmotion analysis; signal type: Dopamine, second order Butterworth low pass filter:10, order 2, peak distance: 100, peak absolute threshold: 2, Time block: 60 seconds.

#### Behavioural test

Open Field test - Distance travelled in the open field (45 x 45 cm arena) was recorded using an acA1300-60gc GigE video camera (Basler). To start the test, mice were released in the corner of the arena facing the wall. Horizontal and vertical activities wwereere recorded for 30 min. For analysis, Ethovision-13 software (Noldus, The Netherlands) was used. Various behavioural parameters such as run straight and turning were scored manually using Ethovision software. All experiments are calculated from 7 mice before and after amphetamine treatment. No mice were removed from the experiment.

#### Amphetamine treatment

Following 30 min open field behavior tests, mice were injected with Amphetamine (5 mg/kg) intraperitoneally and allowed to rest in their home cage for 10 min. Then mice next underwent 30 min open field.

## 3. Results

### 3.1 Testing FPmotion with experimental data

To demonstrate the functionality of the FPmotion software (overviewed in Fig. 1), we used the dLight1.1 reporter to measure extracellular dopamine in the ventral striatum of behaving mice in an aversive space, before and after amphetamine administration. Dopamine release in this brain region is important for initiation of movement in response to a motivational signal (21). We measured peak parameters and dopamine patterns that accompanied behavioral syllables associated with the amphetamine hyperactivity response i.e. turning and running behaviors (Fig. 2A).

**Figure 1.**
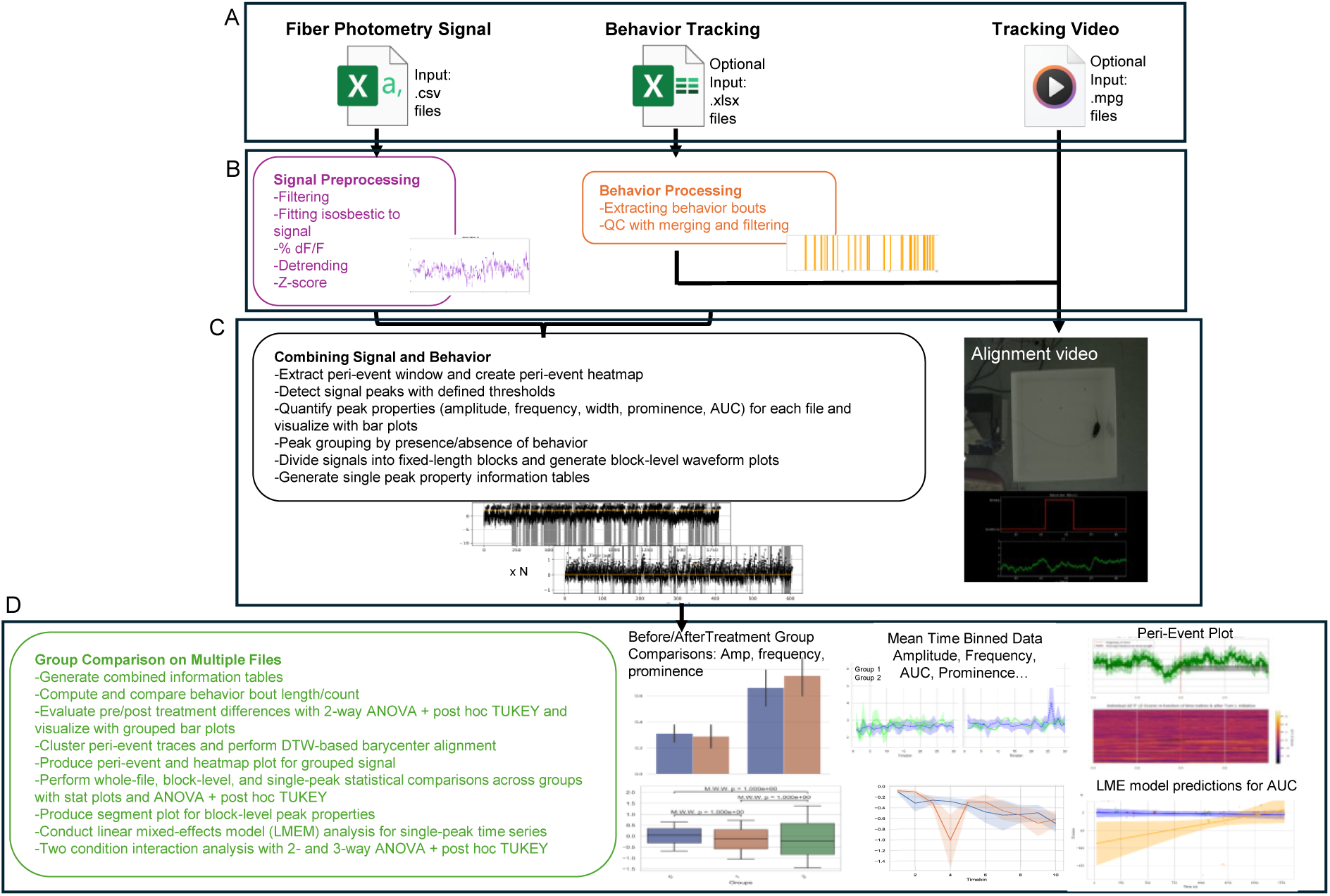
Workflow of the software. **A.** FPmotion software takes fiber photometry signal files and optional behavior tracking and tracking video files, and applies signal processing (low pass filtering, fitting to the reference isosbestic signal, normalisation by calculation of % dF/F, detrending (optional moving average or Butterworth), and z-score calculation. **B, C.** As an option, video files can be imported and combined with the respective signal file and behaviour time-stamped file to generate an alignment video where the fluorescence reporter signal (green) is synchronized with the animal motility in the tracking video and the behaviour bout tracking (red). **D.** Results from all files are then compiled and analysed together with user customized groups and comparisons. The output is organised into a data library (Supplementary data – User Guide Software documentation). Output results include graphical views according to z-score line graphs with and without binning, heat maps, peri-plots and histograms. Statistical test adjusted p-values are illustrated on the graphs as shown, n=7. All data is also output in .csv file with complete statistical breakdown.

**Figure 2.**
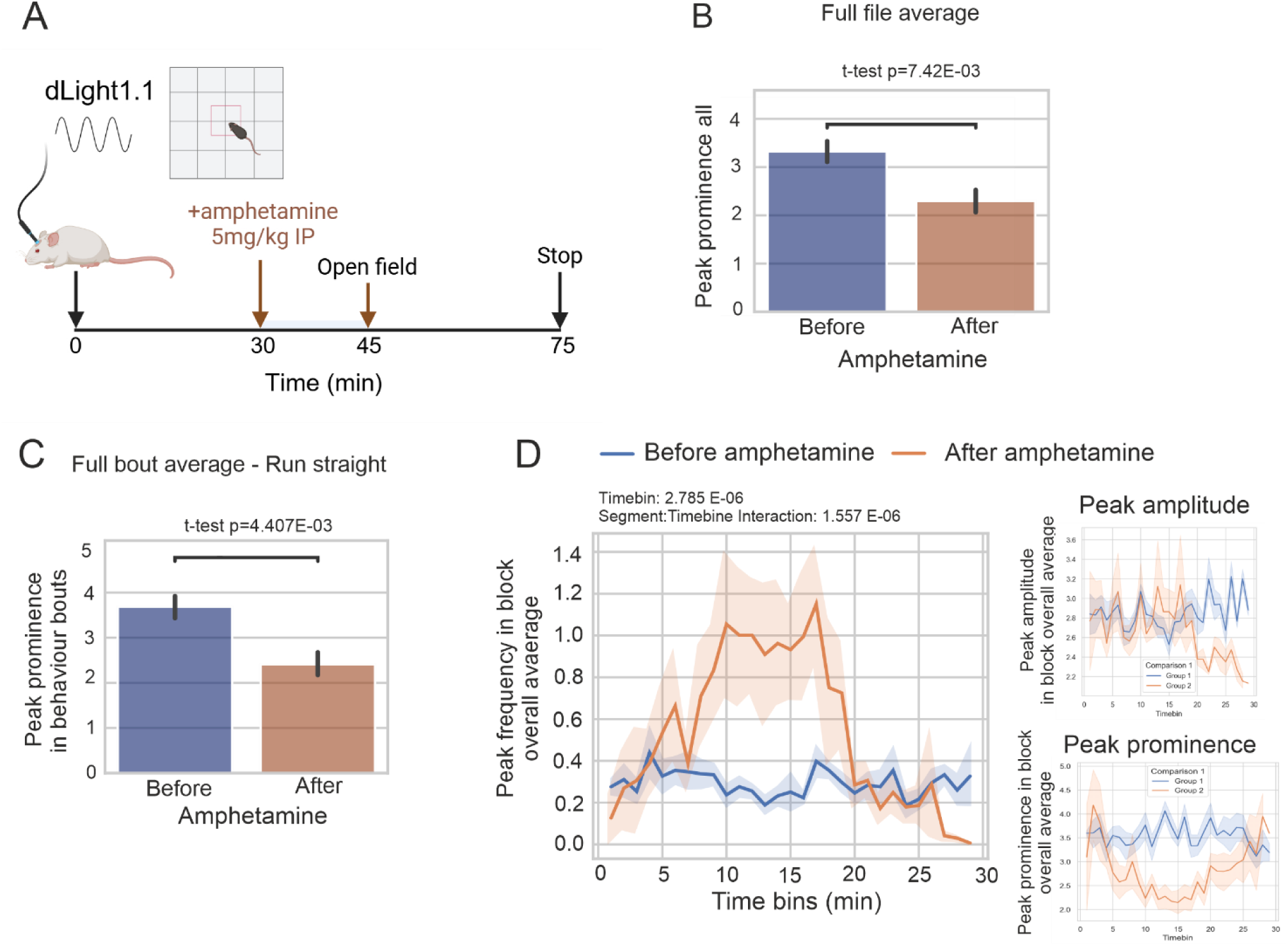
Full file average output before and after amphetamine. **A.** The time course of the mouse experiment is shown. Mice with implanted optic fibres were connected to the fiber photometry console and allowed to move freely in the open field arena for 30 min. Amphetamine or saline was administered I.P. and animals allowed to rest for 15 min after which dLight1.1 signal and behavior were again monitored. **B.** Example output from FPmotion for “peak prominence” based on the total file output for mice before and after amphetamine. **C.** An example of peak prominence output from within a behavioral bout, here shown for “run straight” bouts, before and after amphetamine. Significance for these outputs is determined with Student’s t-test as indicated on the graphic output shown. **D.** Frequency, amplitude and prominence measured from 1min time blocks provide a temporal overview of these parameters before and after amphetamine. Frequency is significantly increased. Significance values for time block data is from two-way ANOVA. All graphical outputs are also generated in csv format. Statistics are shown above the panels. Linear graph p-values are from two-way mixed ANOVA, n=7.

Spontaneous behaviors were captured using Ethovision XT setup and the photometry signal and behavior tracking were saved as .csv files for automated analysis using FPmotion software. The use of FPmotion software to analyse the fiberphotometry and behavior data is described first (2.2), followed by a description of the results obtained in these experiments using FPmotion (2.3).

### 3.2 Software setup

Here we walk the user through the software setup and parameter selection. Data analysis was performed through the FPmotion graphical user interface (GUI). In the Input interface, we loaded the photometry signal files acquired with Doric Lenses FP console and the corresponding behavioral tracking files exported from EthoVision XT. Behavior videos can be uploaded at this stage. Parameter selection was done through the Settings interface. Under “General”, we kept the default settings which skip genotype-treatment combined analyses (as this is used when treatments are done in more than one type of mouse, e.g. in different genotypes). We left the “skip video creation” option at its default disabled state. For “Ethovision” settings, we used FPmotion’s default starting row of 36, the default 7s maximum behavior-bout merging threshold, and the default 1s minimum bout-length threshold. Under “Doric”, we retained the default fluorescence and isosbestic channel labels (“Aln-1” and “Aln-2”). Under “Video”, the frame rate down sampling remained at the default value of 10 fps, although this parameter had no effect because no videos were loaded. For “Peak extraction”, we kept the default setting for minimum inter-peak distance of 100 ms in this instance where acquisition is 1000 Hz) and the default 60s time-block duration for time-resolved analyses. Under “Plot parameters”, all figure types used the default y-axis limits. For “Segment modelling”, we retained the default setting to enable automatic detection and removal of outliers, and we kept the default option that aligns before/after behavior-triggered signal segments as a single continuous trace rather than splitting them into pre- and post-components. We also kept the default option to perform follow-up analyses on barycenter-aligned segments, using the default maximum of 10 clusters, and retained the default minimum peak-distance threshold of 5.

The only modifications we made were in the “Signal preprocessing” and “Peak extraction” settings. In “Signal preprocessing”, we changed the signal type from the calcium preset to dopamine, as dLight 1.1 (and not GCaMP) was the sensor used in these experiments. Accordingly, we increased the frequency threshold to 25 Hz and applied a second-order Butterworth low-pass filter to remove high frequence noise. In “Peak extraction”, we adjusted the amplitude-based peak-detection threshold to an absolute value of 2. Next, we proceeded to the Information Input interface. File metadata were automatically populated from filenames following FPmotion’s naming conventions. After confirming file information, we opened the Group Comparison interface and created a single comparison contrasting the pre-amphetamine (group 1) and post-amphetamine (group 2) recordings. After saving this comparison, we returned to the Information Input interface to complete the remaining configuration, including behavioral selections and z-score normalization.

From the auto-detected EthoVision behavior labels, we selected “Run straight,” “Turn L,” and “Turn R” for inclusion in the analysis. For z-score normalization of the Doric signals, we selected “External” baseline mode and used all pre-injection recordings across mice as the baseline distribution. Because the entire pre-amphetamine window was used, no specific baseline time interval was entered. The user can alternatively select another baseline mode if desired. Genotype color settings were left at their default values. After confirming all parameters across the four interfaces, we initiated the analysis. FPmotion then executed all processing steps autonomously taking 2 hours to complete the analysis from 14 trials (30 min trial duration, 1000 Hz) on a Windows laptop with 32GB RAM. If generation of alignment videos is selected, it takes 4 hours. Upon completion, all graphical outputs, spreadsheets, statistical summaries, and processed signal files were available in the designated output directory.

### 3.3 Sample workflow measuring amphetamine responses with dLight1.1

We measured dopamine signals and behavior from mice freely behaving in an open field arena before and after amphetamine administration. Extracellular dopamine was measured using the dLight1.1 sensor made from a fusion of a D1 receptor fragment with circularly permutated GFP. dLight1.1 is excited by blue light and emits with a peak at around 515 nm. The resulting signal provides sub second time resolution and sub-micromolar sensitivity for dopamine detection, suitable for measuring tonic and phasic dopamine release (4, 22).

Figures 2 to 5 present some of the results from the automatically generated output. Two weeks following viral injection and cannula implantation to the nucleus accumbens (NAc), mice underwent fiber photometry and behavior tracking. Freely behaving mice were monitored during 30 min in the open field after which amphetamine (5mg/kg) was injected intraperitoneally. After 15 min rest, mice were returned to the arena and behavior monitored for a further 30 min (Fig. 2A). Full file averaged data before and after amphetamine shows that peak prominence was significantly reduced after amphetamine whether measured across the entire 30 min monitoring (p=7E-03) (Fig. 2B), or within “run straight” behaving bouts (p=4E-03) (Fig. 2C). Peak prominence is the vertical distance between the peak and the lowest contour line that connects it to higher peaks or the signal baseline. The decrease in peak prominence during this period is likely because the extracellular dopamine is high due to amphetamine inhibition of the dopamine transporter (DAT), preventing a return to baseline dopamine between the peaks. The linear graphs (Fig. 2D), show 1-min time binned output for frequency, amplitude and prominence. Frequency increased sharply 20 min following amphetamine injection and remained elevated for 15 min after which the plateau diminished. The high frequency period showed good overlap with reduced peak prominence period (Fig. 2D), consistent with dopamine being highly elevated. Amplitude was not significantly altered by amphetamine when looking at the full file output, although it visibly decreased at the point when frequency returned to baseline. These data closely match the timeline of dopamine release measured by microdialysis following intraperitoneal injection of amphetamine (23).

FPmotion also analyses single peaks. Figure 3 shows a selection of outputs from the “single peak” analysis. Amphetamine decreases prominence once more as shown by plot and scatterplot (Fig. 3A, B). The statistical significance for single peak output is based on LMEM, accounting for within subject variability and repeated measures. Within and outside of behaving windows calculations are automatically generated for all “selected” behaviors within the software. Here we show peak prominence during “run straight” bouts also decreases after amphetamine treatment, visualised by barplot and z-score scatter plot (Fig. 3 C, D).

**Figure 3.**
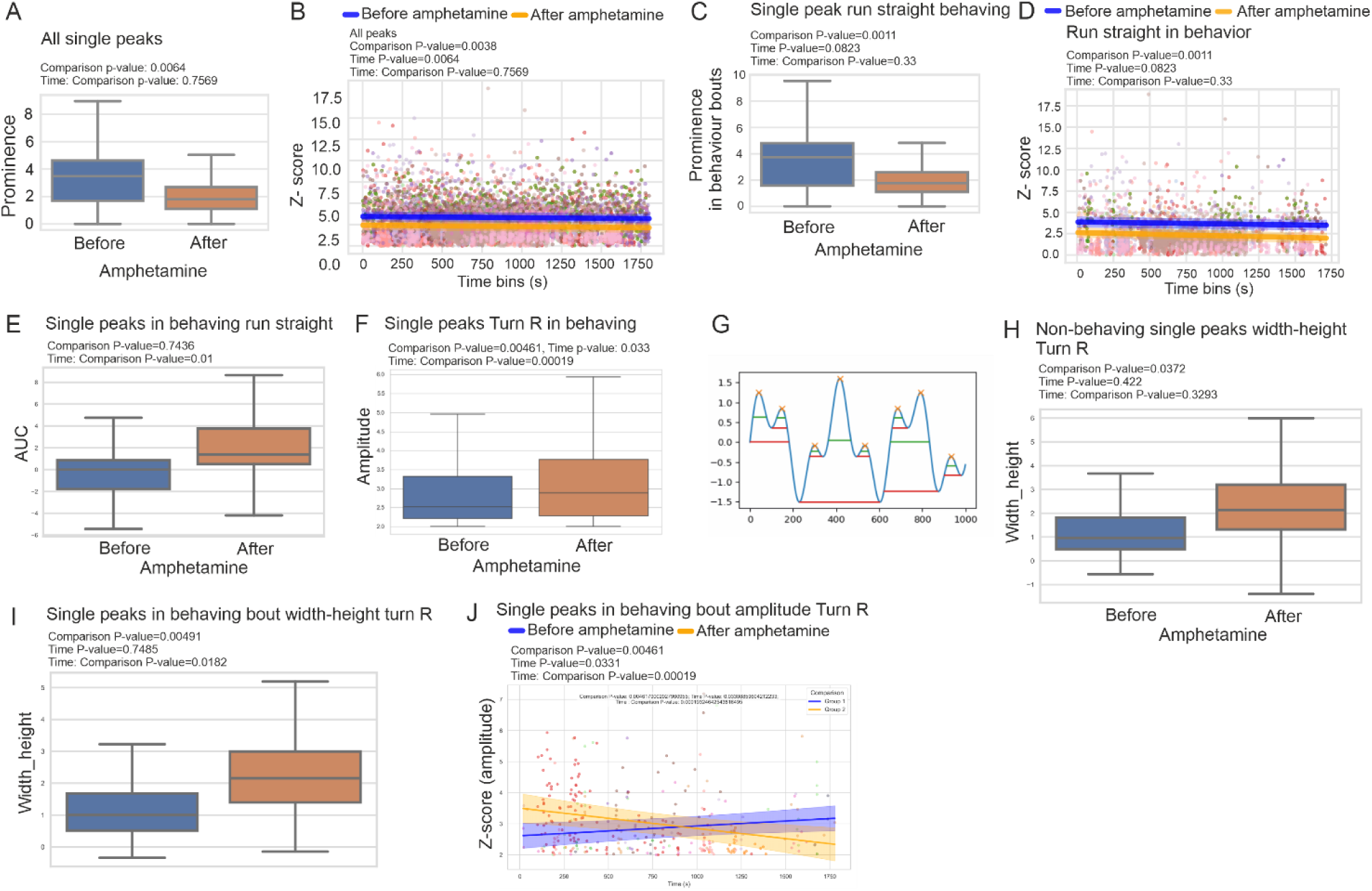
A. Single peak output before and after amphetamine. **A.** Single peak data for peak prominence before and after amphetamine is shown in a bar plot and **B,** as a scatter plot (z-score). Data is calculated from single peaks before and after amphetamine administration. Statistics are from LMEM. **C, D.** Single peak data for prominence during “run straight” in behaving window is shown as bar plot and scatterplot, before and after amphetamine. **E.** A.U.C of peaks during “run straight” is shown. **F.** Single peak amplitude before and after amphetamine during “turn right” behavior bouts. **G.** A scheme illustrating peak width-height. **H, I.** Peak width-height results during “turn right” bouts is shown before and after amphetamine in non-behaving and behaving windows respectively. **J.** A scatter plot of amplitude before and after amphetamine in “turn right” bouts. Statistics are from LMEM, (n= 7), Area under the curve (AUC) within “run straight” bouts is significantly changed with respect to time (Time:comparison p-value = 0.01) (Fig. 3E). Although there is no significant difference between the groups for the average overall AUC values, there is a significant change in AUC between the groups with respect to time (p=0.01) (Fig. 3E). We next show an example of amplitude data. Within “Turn right” bouts, peak amplitude increases by 14% (p-value=0.00461) following amphetamine. Amplitude also changes significantly with respect to time (time: Comparison p-value = 0.00019; Fig. 3F).

FPmotion also calculates “peak-width-height”, which is the height of the contours at the point where widths were calculated (Fig. 3G). Peak-width-height increases two-fold following amphetamine. The “turn right” width-height results from non-behaving (p=0.0372) and behaving (p=0.00491) bouts is shown (Fig. 3H, I). A temporal view of amplitude changes before and after amphetamine highlights a significant interaction between amplitude and time, where amplitude is higher after amphetamine but decreases by about 30 min following intraperitoneal injection of amphetamine. This decrease may reflect vesicular dopamine store depletion following amphetamine (24) (Fig. 3J).

Figure 4 shows examples of FPmotion’s calculated behavioral bout output. Following amphetamine, “run straight” bout length increases (p=1-26E-07) (Fig. 4A), while bout number decreases, reflecting prolonged hyperlocomotion following amphetamine (Fig. 4B) (25). For “turn left” syllables, bout length also increases significantly (p=3.08E-05), consistent with amphetamine-induced circling (Fig. 4C) (25). In this case, bout count similarly increases (p=0.00623; Fig. 4D). In contrast, “turn right” syllables show a significant increase in bout length, but no change in bout number (Fig. 4E, F). Individual dopamine responses can be examined from the .csv files and from the synchronized alignment video (Fig. 1C; supplementary video).

**Figure 4.**
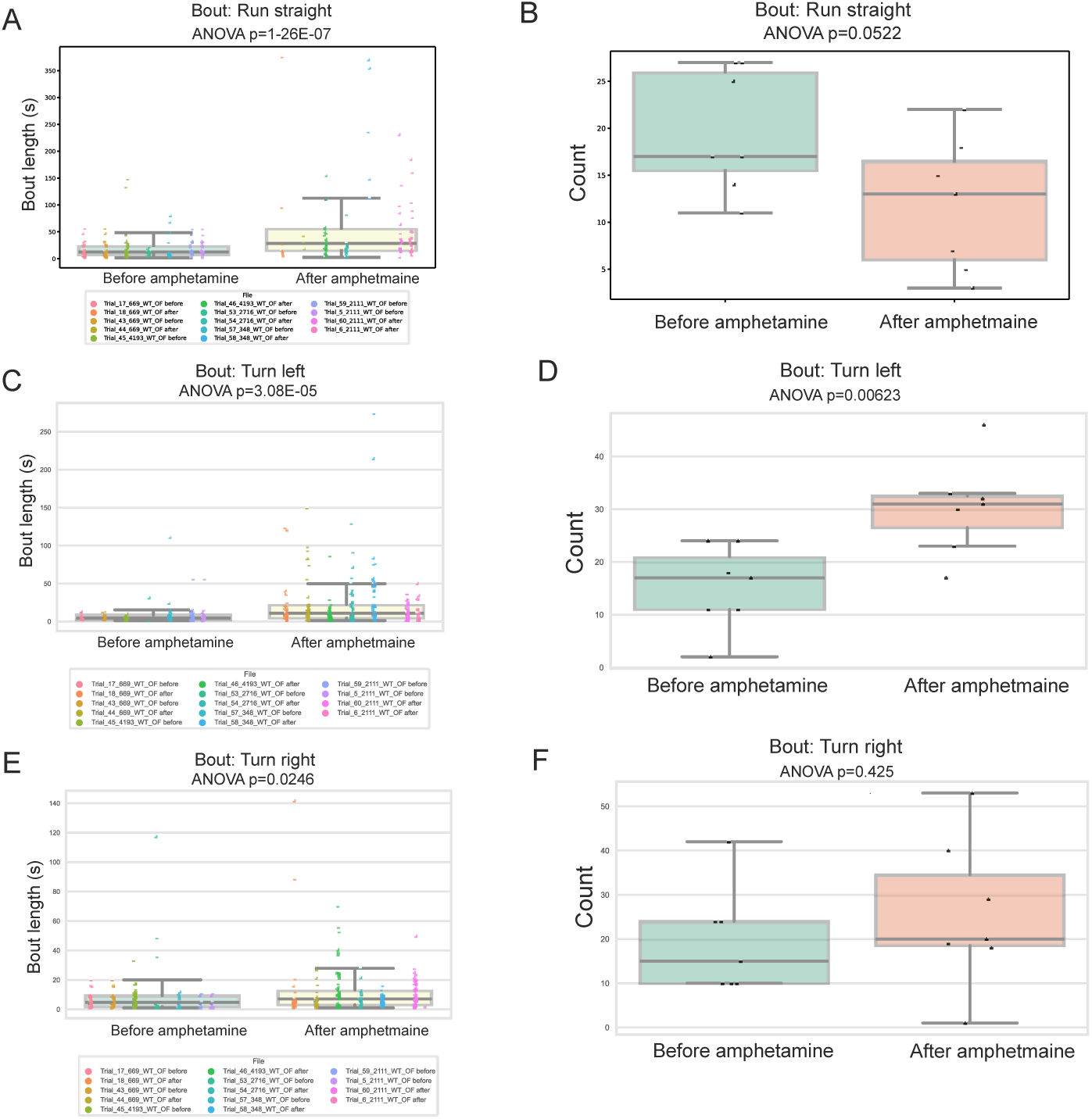
Behavioral output before and after amphetamine. **A, B. “**Run straight” bout length and count are shown before and after amphetamine administration. **C, D.** “Turn left” bout length and count are shown before and after amphetamine. **E, F.** “Turn right” bout length and count are shown before and after amphetamine. Statistical analysis is with ANOVA, n=7.

The peak alignment function is illustrated for 5s before and after behavior initiation as peri-plots (Fig. 5 A, B). The purpose of the peri-plot is to show how dopamine peaks occur relative to behavior bout onset, allowing identification of co-occurring dopamine patterns. Temporal alignment of peaks is necessary to correct for peak latency. This improves signal clarity and reduces errors caused by misaligned signals (Fig. 5B). The outlier function is demonstrated in Figure 5C. Here FPmotion identifies an outlier based on gross difference in amplitude (red). This trace will be removed from the quantification if the “*outlier removal*” option is selected. The user can also choose not to remove outliers. All outliers are highlighted as shown, in the graphical output so that the user can check that they agree.

**Figure 5.**
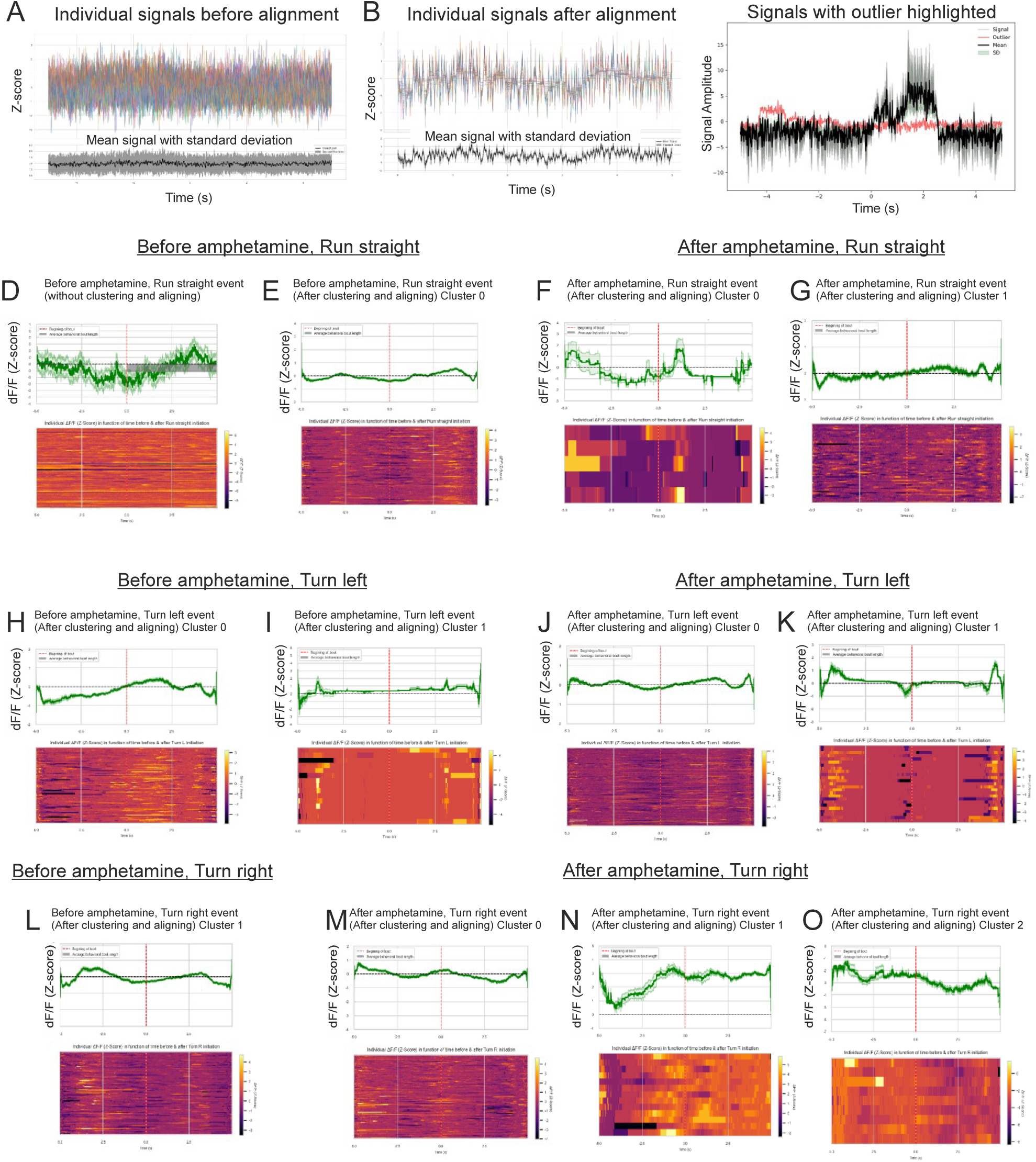
**A, B.** Individual and meaned dLight1.1 signal before and after DTW alignment is shown. **C.** An example of a removed outlier (red) based on amplitude measurements. The mean trace is shown in black. **D.** Before alignment peri-event-plot of mice without amphetamine shows 5s before and after time anchored “run straight” bouts. The lack of alignment is visible from the heat map which shows all events. The dopamine signal is dark green and standard errors are pale green. **E**. Peri-plot of “run straight” events after alignment is shown for mice without amphetamine plotted. **F, G.** Alternative (cluster 0, cluster 1) peri-plots for “run straight” events after amphetamine. The user must use discretion in selecting clusters. In this case, cluster 0 may be an outlier, as there are few occurrences compared to “F”. **H-K.** Peri-plots during “turn left” syllables before and after amphetamine are shown. In each case two alternative clusters are shown. **L.** Peri-plots before, and (**M-O**) after amphetamine during “turn right” syllables, are shown. Three alternative clusters are shown after amphetamine. The similarity in dopamine peaks and troughs suggest that all clusters may represent real bout-associated signals.

Figure 5D shows “run straight” events *before* dynamic time warping-based peak alignment and clustering. The signal to noise ratio is poor and no clear pattern is visible from the heatmap. In contrast, after aligning and clustering, a distinct pattern emerges (Fig. 5E). This shows that dF/F (z-score) dips before initiation of “run straight” followed by a gradual incline (Fig. 5E). Clusters 0 and 1 after amphetamine represent possible dopamine patterns (Fig. 5F, G). dF/F (z-score) for cluster 1 (Fig 5G) resembles the “run straight” pattern before amphetamine (Fig. 5E). However, as cluster 0 for “run straight” after amphetamine (Fig. 5F) only occurs 5 times, this signal is likely the result of noise and would be considered an outlier. Note, the cluster numbering is arbitrary and does not signify rank. Also, the user should use discretion in deciding which clusters are likely to represent authentic signals.

Turning behavior is associated with asymmetry in dopamine signaling, where turn direction is towards the hemisphere with lower dopamine levels (26), while bilateral dopamine elevation produces generalized hyperlocomotion, as reflected in this case by increased run straight bout length (27) (Fig. 4). Note that in these experiments, the fiber photometry canula is implanted in the right hemisphere. Consistent with this, the peri-plots for “turn left” syllables before and after amphetamine show increased dopamine when turning initiates (Fig. 5H, J). However, another pattern is also seen during turn left where phasic dopamine release is observed -2.5s before turn initiation and +2.5s after (Fig. 5I). Following amphetamine, the same peaks are observed (Fig. 5K). While the pattern is identical before and after amphetamine, a greater proportion of “turn left” bouts display it (15 bouts before verses 25 bouts after, I and K), suggesting that it represents an authentic dopamine signal ensemble.

We also examined peri-plots for “turn right” behavior syllables. Dopamine in the right hemisphere dipped before “turn right” behavior commenced in control trials (Fig. 5L, cluster 1), however, after amphetamine administration, three signal ensembles emerged from the clustering analysis (Fig. 5M-O). All clusters showed a dip in DF/F (z-score) from -5 to -2 s before turn initiation at timepoint 0, by which point dopamine was higher. This was followed by a dip in dF/F (z-sc) signal following turn right initiation (Fig. 5M-O). Given the reasonable signal to noise ratio and the repeated nature of these ensembles, we propose that they represent bonafide NAc dopamine patterns accompanying turning. These data show that FPmotion peri-event plots successfully resolve discreet ensembles within the data.

## 4. Discussion

FPmotion balances a structured analytical workflow with the flexibility needed to accommodate diverse experimental designs. The core processing steps, spanning signal preprocessing, behavior-event extraction, normalization, visualization, and advanced analyses, is organized to allow users to obtain meaningful results from a single analysis. At the same time, most steps of the workflow include adjustable parameters, allowing the analysis to be refined for example for different biosensors. Although the software provides default parameter choices that are robust, users can adjust filtering cutoffs, normalization modes, and event-alignment windows. Group comparisons can also be configured. In this way, the framework provides consistency for users preferring a turnkey approach while remaining adaptable.

The software generates a comprehensive set of results during the analysis run, rather than requiring users to preselect specific outputs. This approach minimizes the risk of missing relevant summaries, and the need to repeat the analysis. By providing all outputs at once, researchers can focus on the figures and tables most meaningful for their experimental questions. These files can be easily viewed in the FPviewer software. This analysis environment is easy to navigate and supports a broad range of users.

The final and most critical component of the workflow is the alignment of peri-event signals using DTW barycenter averaging (DBA). Behavioral onset alone does not guarantee temporal alignment of fluorescence features across trials; even after time-locking all segments to the same event. The latency of prominent peaks and valleys in FP traces varies between trials. Such latency jitter is well documented in other neural modalities, such as event-related potentials and calcium imaging, where a simple stimulus-locked averaging attenuates waveforms unless temporal realignment is applied (28, 29). To address this, we incorporated DTW-based alignment, constrained by the fixed behavioral time anchor, permitting local latency corrections while preserving the event reference. We adopted DTW DBA to compute a consensus waveform for each cluster. DBA provides a more faithful representation of average temporal shape than arithmetic mean, particularly when peaks and troughs occur with slightly different latencies across trials. Prior work highlights its ability to preserve shared temporal motifs in time-series datasets subject to latency variability and non-uniform warping (20, 30). In practice, this alignment pipeline enabled clearer peri-event motifs to emerge that were largely invisible in the raw, unaligned averages despite time locking. Nonetheless, we emphasize that this method is optimal only when the trials share broadly consistent temporal dynamics. Our simulation experiments reveal that alignment performance degrades when large frequency differences occur across signals. Users should therefore use it cautiously when underlying kinetics are highly heterogenous.

Throughout the development of this workflow, we evaluated method performance using simulated datasets that varied in amplitude, asymmetry, frequency, and polynomial shape. These tests revealed that the pipeline is sensitive to data characteristics, particularly signal frequency (Supplementary data 1). DTW-based approaches perform well when signals share broadly similar dominant frequencies, as is typical for FP dopamine or calcium recordings. DTW may attempt to warp high-frequency structures into low-frequency ones, producing biologically unrealistic alignments. This underscores the importance of empirically validating outlier removal, clustering, and alignment decisions, and avoiding overinterpretation of motifs that emerge from datasets with high-frequency variability or poor signal-to-noise ratios.

In summary, the combination of PCA-based Mahalanobis distance calculation, variance-weighted MCD estimation, fast DTW clustering, and DBA-derived consensus templates alignment provides a robust and scalable framework for detecting meaningful temporal patterns in peri-event FP signals. The pipeline is specifically designed for datasets with many trials, moderate event-related variability, and relatively consistent signal frequency; conditions typically met in dopamine and calcium FP experiments. Overall, the methods implemented here provide a principled and computationally fast and automated way to uncover reproducible FP dynamics surrounding behavior with batch processing, full statistical analysis and graphical, .csv and video output.

We tested the pipeline with fiber photometry recordings from the ventral striatum of mice before and after amphetamine to investigate what transient dopamine signals accompanied behavior syllables in freely behaving mice. Our results show that amphetamine increases the frequency of dopamine transients while reducing peak prominence. This makes sense given the sustained elevation in extracellular dopamine upon amphetamine-mediated DAT inhibition. Normally, extracellular dopamine in striatum clears within 100 to 300 ms (31, 32). Following systemic amphetamine injection, we find that dopamine is elevated in ventral striatum by 20 min post injection, consistent with previous reports for amphetamine and cocaine (23, 33). Frequency remains elevated for 15 min from this time point while amplitude and AUC increase significantly. Moreover, peak prominence decreases while peak-width-height increases significantly following amphetamine treatment, consistent with prolonged dopamine availability in the extracellular space. Temporal views of the data reveal changes over time, for example amplitude decreases by 30 min following amphetamine, suggesting a decline in dopamine, and consistent with this, frequency decreases in parallel. Finally, the peri-event plots capture an expected dip in dF/F (z-score) for dopamine in the right hemisphere of NAc when initiating right turns as previously described (33). This improved peak alignment combined with clustering allows detection of discreet signal ensembles that would otherwise be masked. Overall, we demonstrate that the FPmotion features including automation of fiber photometry data processing, outlier removal, modelling, peri-event analysis (with dynamic time warping and clustering) and statistical analysis simplifies the translation of fiber photometry signals into biological insight and makes it accessible to non-programmers.

