## Supplementary data for "FPmotion an Automated Signal Processing and Statistical Analysis Tool for Fiber Photometry Data"

A

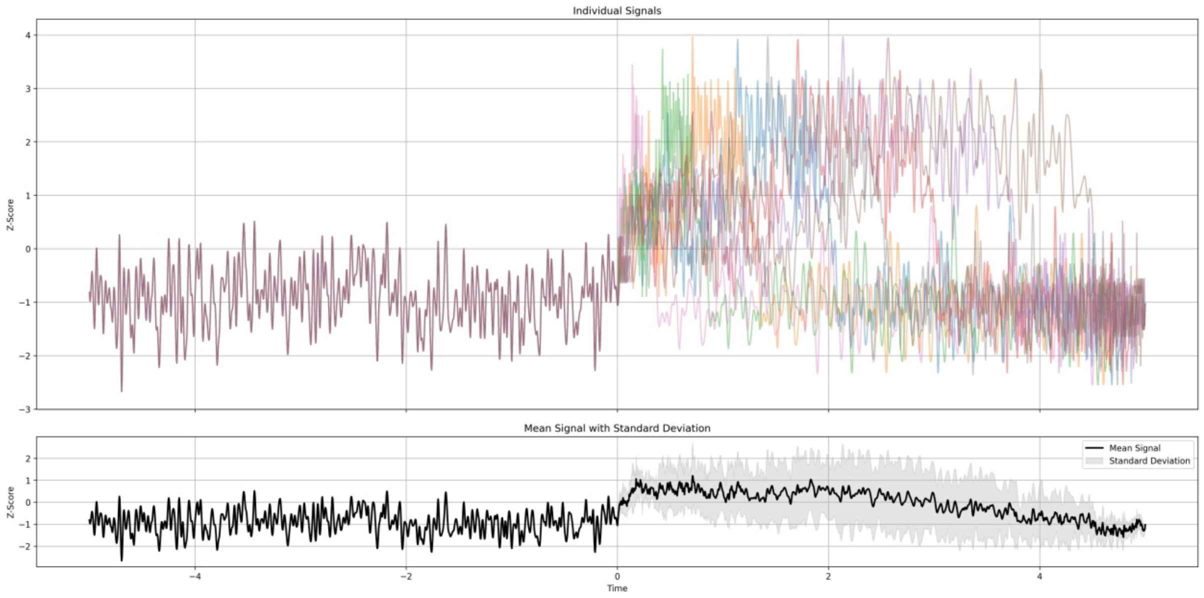


B

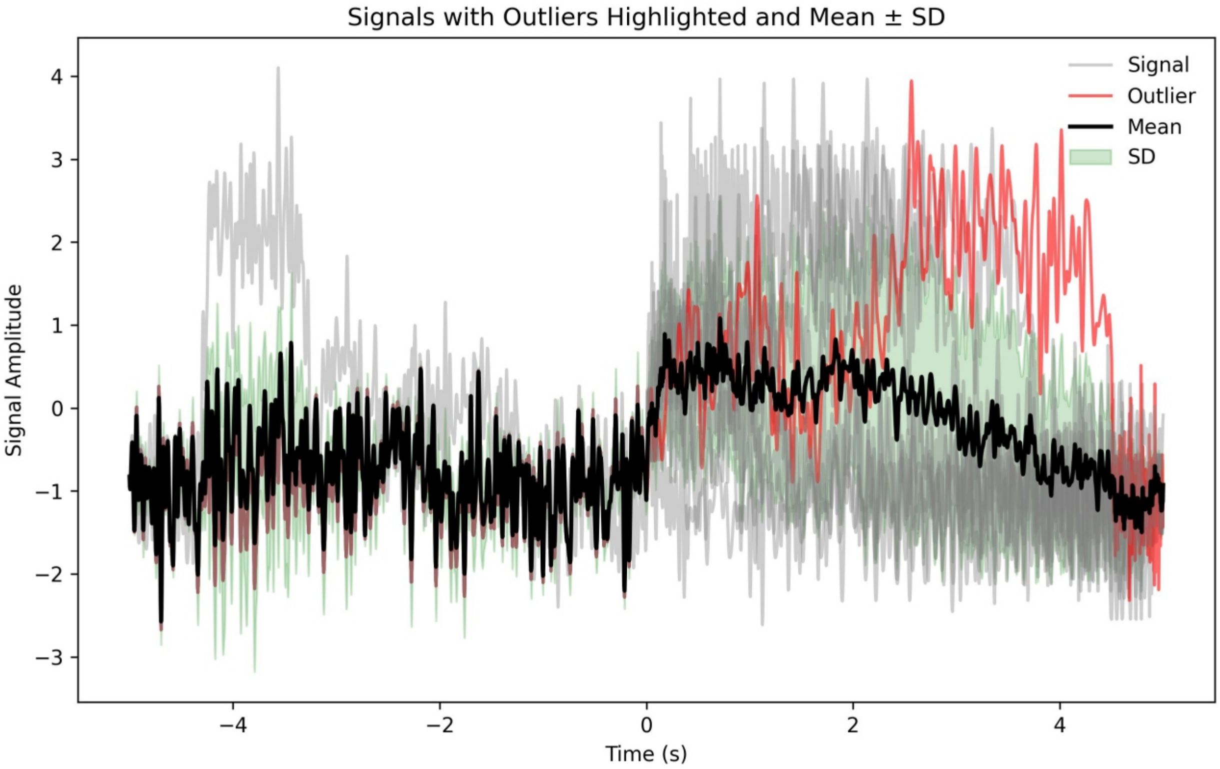


**Supplementary Data 2**. Identification and removal of outlier based on frequency difference. A. Simulated data with different frequencies are shown including the mean trace. B. When outlier removal is selected, the red trace is identified as an outlier based on it having lower frequency than the other traces. The mean (black trace) excludes the outlier. Users should carefully check outliers if this optional step is utilized.

By Ye Hong, Jismi John from Coffey Lab

### FPmotion User Guide

### Input Interface

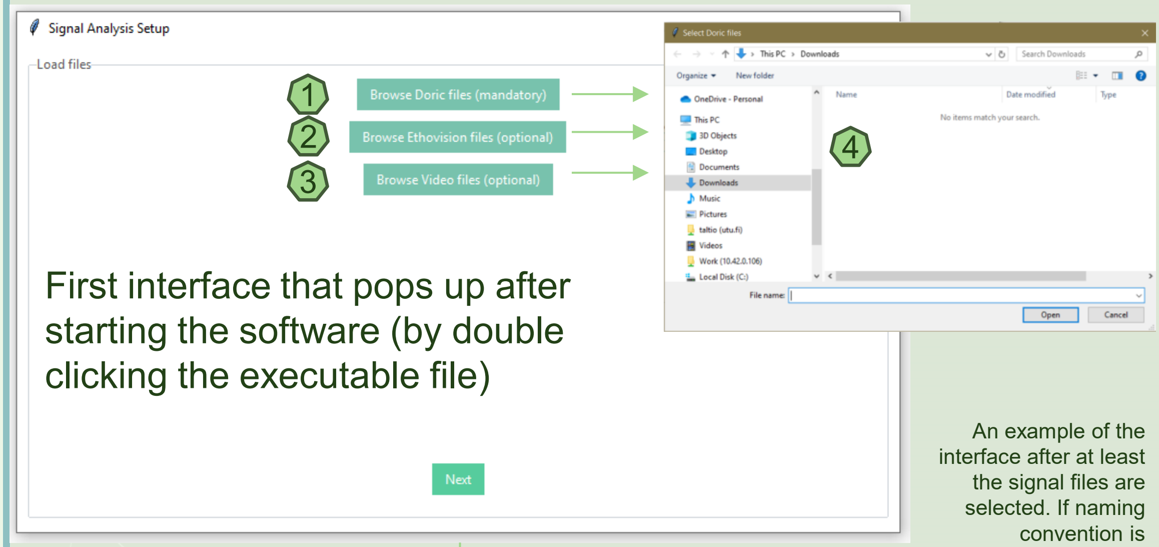


First interface that pops up after starting the software (by double clicking the executable file)

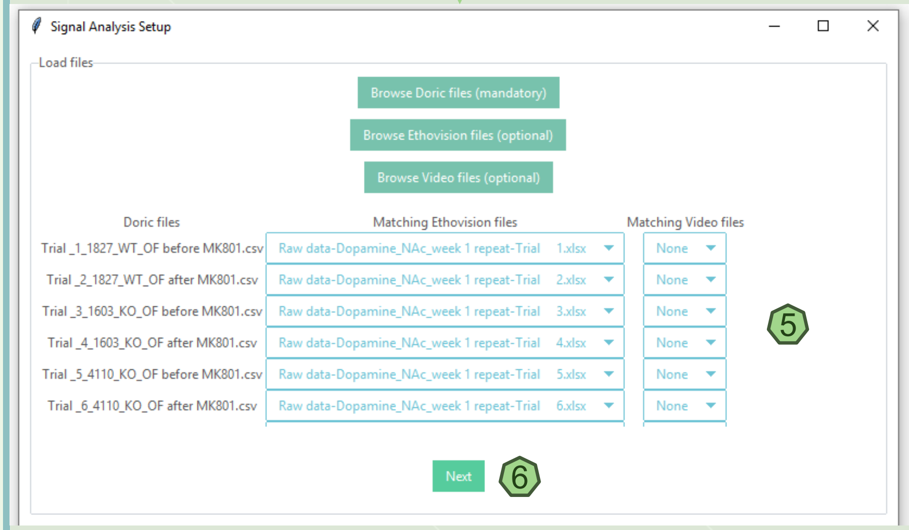


5 An example of the interface after at least the signal files are selected. If naming convention is followed, the signal files will be put in order of the trial number, then the tracking and tracking video files will be ordered by matching signal file's trial number. For tracking and video files, clicking on the file name opens a dropdown menu where all selected files are in the list and a different file can be selected.

6 After the setup is done, click on Next to bring up the parameter interface.

1 Click to select fiber photometry signal input files. The default file is signal output from Doric equipment; however, all files can be accepted with the following format: CSV format where column 1 records time points, column 2 records isosbestic channel, and column 3 records dopamine/calcium channel. See Parameter Interface Number 4 to indicate columns if default column name is not AIn-1 and AIn-2.

2 Click to select behavior tracking input files. The default input is tracking file generated by Ethovision software; however, any tracking files can be accepted with the following format: Excel format where "Recording time" column records time points, and other columns record behaviors, one per column. Behavior columns should have values 0 or 1, where 0 indicates behavior not present at the matching time point, while 1 indicates behavior is present. Change "File starting row" in Parameter Interface Number 3 to 1. See Parameter Interface Number 3 for other behavior processing settings. This input is optional.

3 Click to select tracking video files. Any format supported by FFmpeg can be accepted. This input is optional.

4 An example browse window that will open after clicking 1-3. Multiple files from the same folder can be selected. Select files as you would with a standard file explorer, ctrl+click, shift+click, ctrl+a, and mouse highlighting all apply. Click on Open when the files are selected.

### Parameter Interface: I

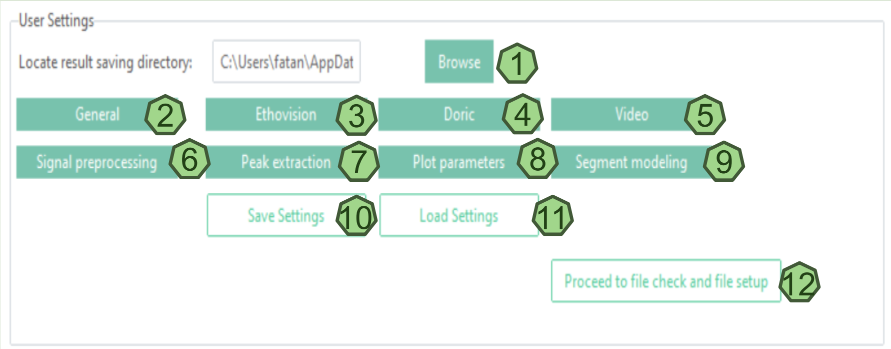


1 Click to select the result saving directory.

2 Click the general tab to perform optional tasks (see 2.1). The first option could enable two-conditions analysis, and the second option determines whether the output includes combined video results

3 Click the Ethovision tab to set file parameters such as the file starting row, minimal bout length and behaviour merging distance (See 3.1). (File starting row means the row on the behaviours excel where the behaviour and timestamps headings are mentioned. Some Ethovision files have the first 35 rows where the experiment, animal numbers, etc. is described and the information headings such as start time, run, entries into arms, etc. are mentioned only from row 36 onward.) The user can define the distance between 2 different adjacent bouts, and the ones that are closer than the defined distance will be merged and considered as s single bout. The user can also define the minimum duration of a bout, shorter than the user defined minimum duration will be excluded.

4 Click the Doric tab to change the signal and background channel names according to the csv files you have (See 4.1).

5 Under the video tab, you can downsize the video frame rate, which could decrease the runtime of the combined-video generation (See 5.1).

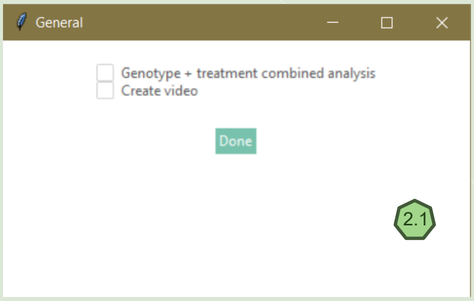


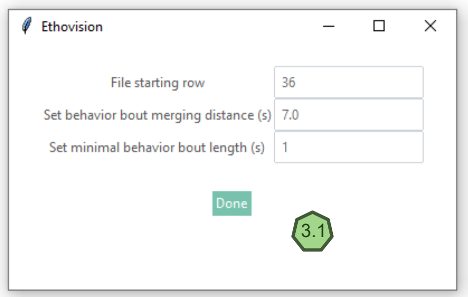


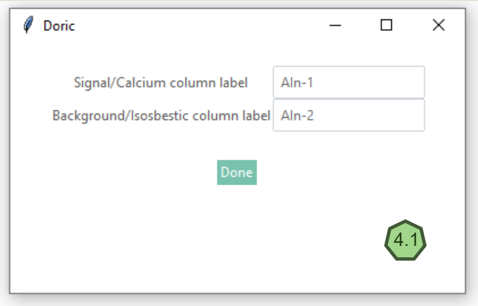


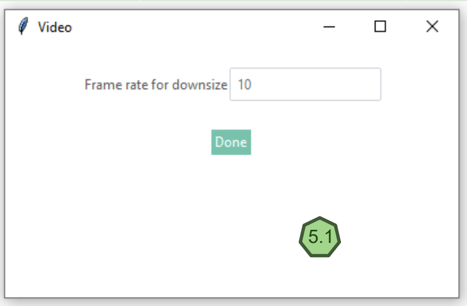

### Parameter Interface: II

6 Signal pre-processing tab: The user can select between 2 types of signal processing, “Calcium” or “Dopamine”.

- Calcium signal pre-processing includes a zero-phase moving average digital filter applied backwards and forwards. The user can set the filter window size in the unit of data point counts (See 6.1).
- Dopamine signal pre-processing includes a second order Butterworth low pass filter. The user can set the frequency threshold (Hz) and the order of the low pass filter (See 6.2).

7 Peak extraction tab: Here the user can set the peak distance, time length for block analysis (in seconds) as well as the type of calculation for peak threshold, relative (MAD*2.91) or absolute threshold value (See 7.1 and 7.2). Peak distance corresponds to the rise time and the half life (1/2) of the reporter used and any peaks lower than that will be considered noise. Time length of the block analysis corresponds to the duration of the  time binned segments for visualising the data in a linear graph. Peak thresholds define signal peaks, with customizable relative and absolute thresholds. The relative threshold uses the median absolute deviation of the signal z-scores multiplied by 2.91, while the absolute threshold is user-defined.

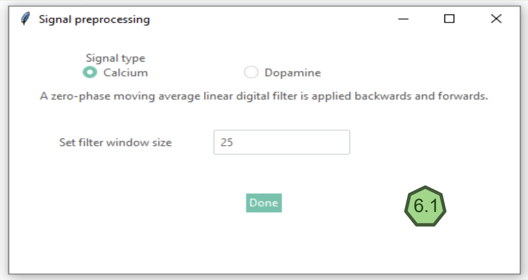


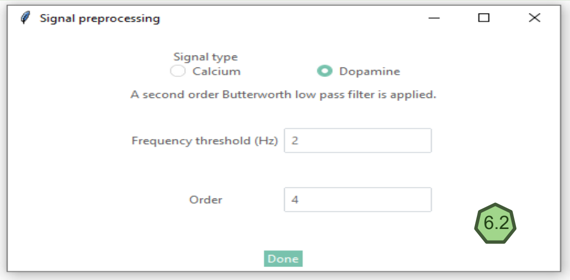


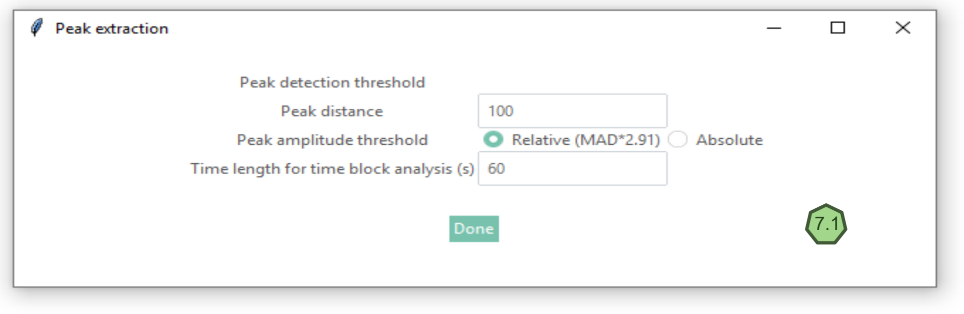


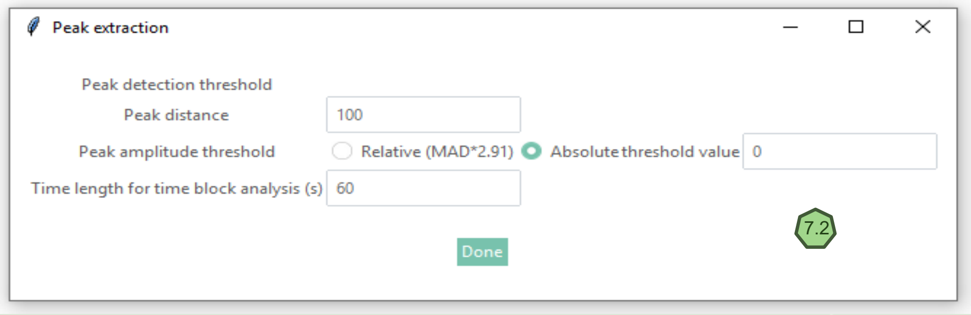

### Parameter Interface: III

8 Plot parameters: In this tab you can specify x- and y- axis range and other settings for specific figure types including general plots, time bin plots, heatmaps and single graph/ figure traces. The default option will configure the best windows for each individual figure separately, making comparing figures difficult. If you would like to, for example, compare the same figure for different files, you can specify the figure parameters for the figure type so they would have the same setting ranges and be easily comparable (See 8.1).

9 Segment modeling: This tab includes modeling parameters for before/after bout segment alignment processing steps. The user can decide whether to include remove outlier signal step as part of the processing pipeline with the first check box. With the second check box, the user can decide whether the before/after signal combination should be aligned together or aligned separately then put together subsequently. With the third check box, the user can decide whether to perform peak analysis for the aligned signals. The user can input a maximum number of clusters for the clustering step in the first input box, where the optimal cluster number will be determined from one to the maximum number of clusters the user chose. In the second input box, the user can determine the peak distance threshold for the follow up peak property analysis.

10 11 User can save the above settings as well as use the settings that were saved from a previous experiment.

12 Click on “proceed to file check and file set up” to bring up the next setup window: File check and comparison setup.

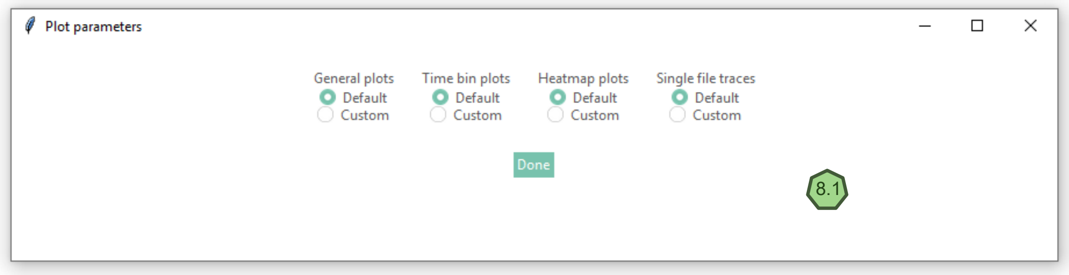


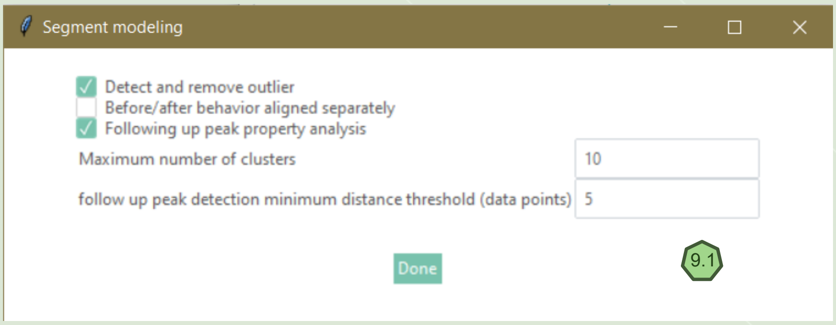

### Information Interface: I

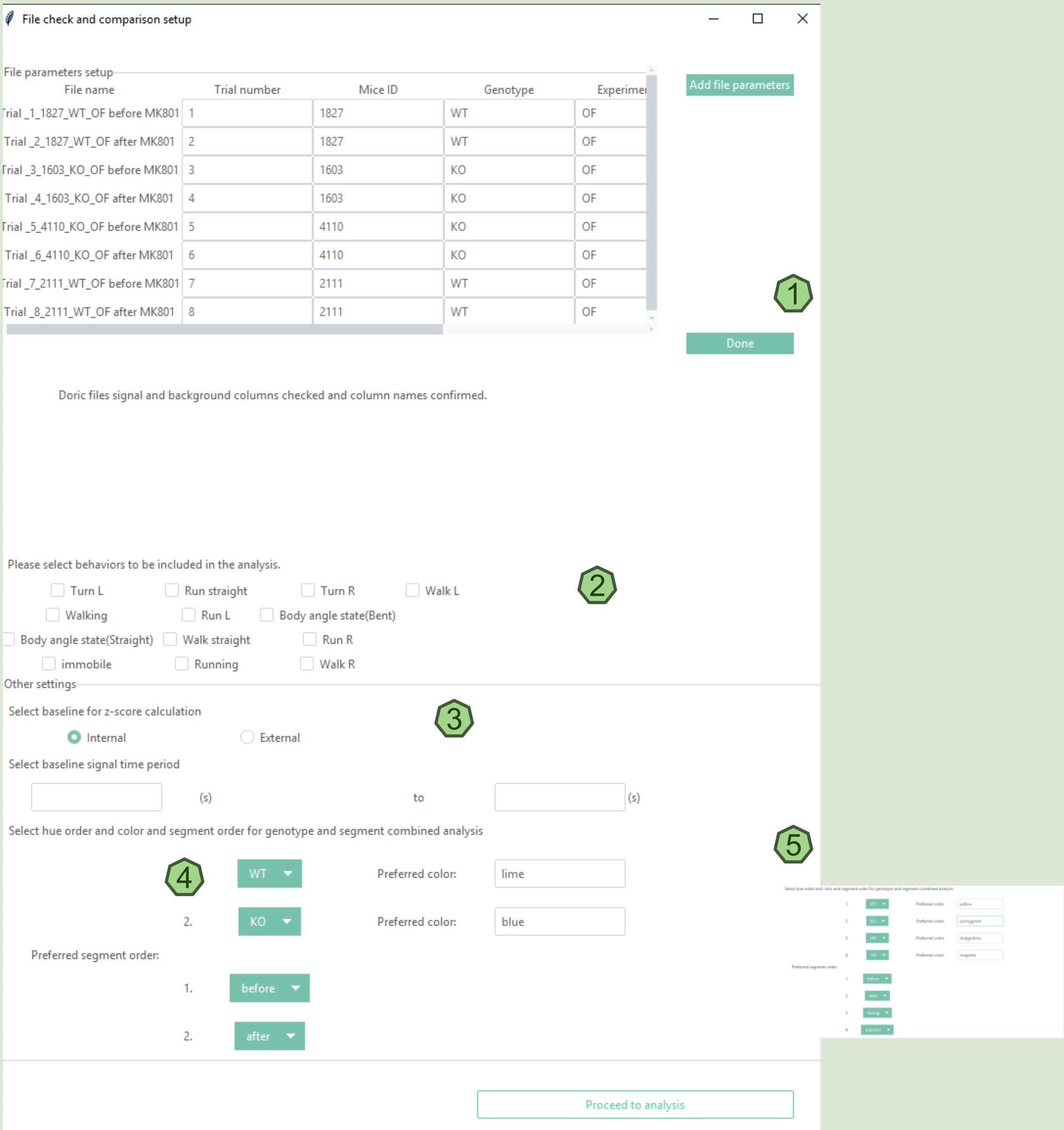


1 Once the software checks the files, a new window named “File check and comparison set up” opens up (Step 3). In this window, the user can add parameters such as trail number, Mice ID, Genotype, experiment type, segment and treatment. If the input files follow the naming convention on page 8, these information should be pre-filled. There are provisions to add more parameters too. Once the file parameters are set up, click “Done” and a new window named “ Group comparison set up” opens.

2 The next step is to select the behaviours to be included in the analysis. All the behaviours that are present in the Ethovision behaviour files will be shown here. Multiple behaviours can be selected/checked.

3 The baseline for z-score calculations can be done in two ways: Internal and external (more information on the next page).

4 In the next step, the user can select colour and segment order for genotype and combined analysis. Proceed to analysis. The user can define any number of variables in genotypes and segments to be included in the analysis (see 5).

### Information Interface: II

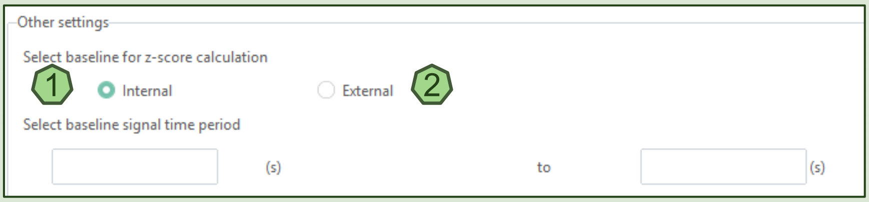


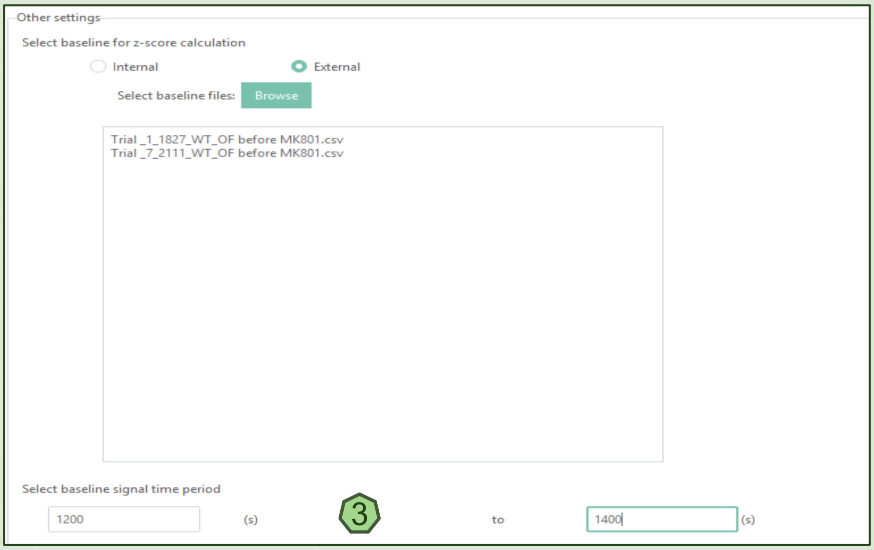


The baseline for z-score calculations can be done in two ways: Internal and external.

1 In the internal baseline calculations, baseline is calculated within each signal file.

2 In the external baseline calculations, the user can select the files that has to be used as a baseline for the signal processing.

3 It’s possible to use only part of the full signal from the selected external files for baseline. The user can specify the part of the signal to use in seconds from these files. Note this is not available for internal baseline calculations.

### Group Comparison Interface

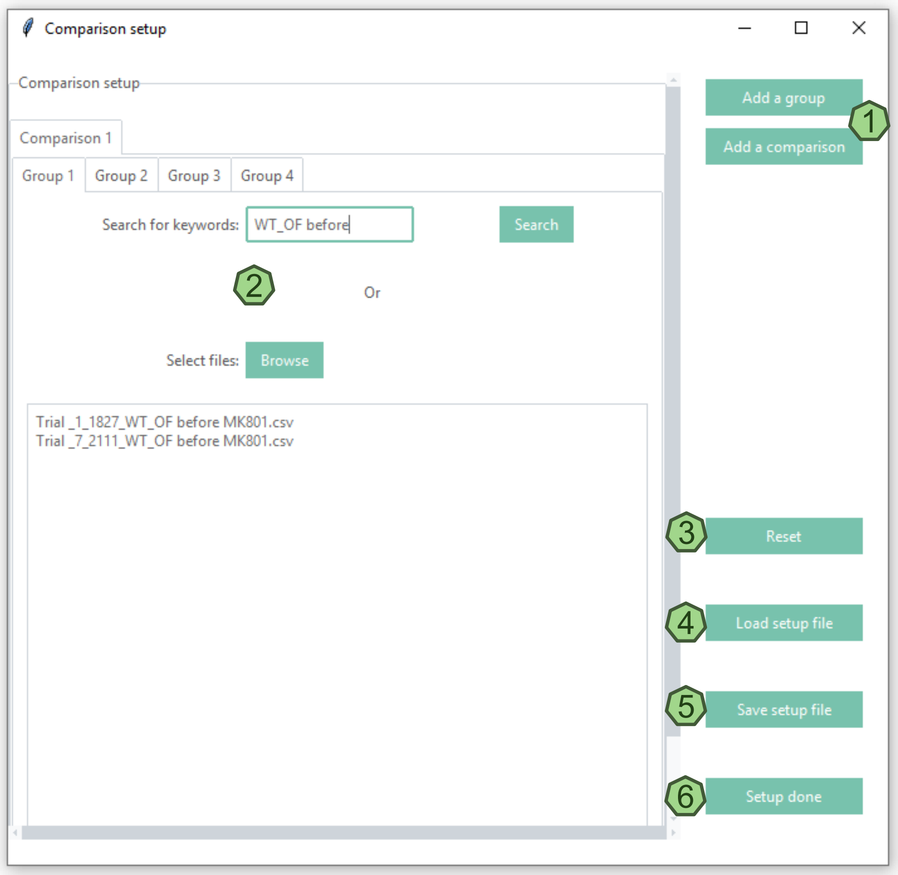


1 Here the user can add multiple comparisons as well as add multiple groups in each comparison. Clicking "Add a group" will add a group tab (Next to "group 4" in the visual); and clicking "Add a comparison" will add a comparison (Next to "Comparison 1" in the visual).

2 The user can add files to each of the groups by searching for it with specific keywords from the file name or by browsing and selecting the appropriate files.

3 The user can reset the comparison by clicking on "Reset". This will erase all the setup in this window. If there was a mistake, e.g. one too many comparisons have been added, "Reset" button can be utilized.

4 5 The comparison set up can also be saved and loaded for future experiments.

6 Once the group comparison setup is done, click on "Setup done" to close the window and incorporate the settings.

### Naming Convention

- Trial _1_1827_WT_OF before MK801
- Trial _2_1827_WT_OF after MK801
- Trial _3_1603_KO_OF before MK801
- Trial _4_1603_KO_OF after MK801
- Trial _5_4110_KO_OF before MK801
- Trial _6_4110_KO_OF after MK801
- Trial _7_2111_WT_OF before MK801
- Trial _8_2111_WT_OF after MK801

1

1

Example input signal file names. To match signal files with their respective behavior files and extract information directly from file names, we implemented specific naming conventions. The signal file name consists of six information fragments connected by underscores or spaces: “Trial_X” (trial number), mouse ID, genotype, behavior test name, and two experiment conditions. For example, "Trial_1_1827_control_OF_before_MK801" indicates treatment ("MK801") and time segment condition ("before"). Behavior and video files only need to match the trial number with the signal files. The trail numbers for the behaviour files and the video files have to be added at the end of the name of the files, eg: “Behvaiour file Trial 1”. Manual adjustments are possible if naming conventions are not followed.

### Result file organization

Name

- Combined information and comparison results
- File 1 Trial_5_2111_WT_OF before
- File 2 Trial_6_2111_WT_OF after
- File 3 Trial_7_4110_KO_OF before
- File 4 Trial_8_4110_KO_OF after
- File 5 Trial_11_12462_KO_OF before
- File 6 Trial_12_12462_KO_OF after
- File 7 Trial_17_669_WT_OF before
- File 8 Trial_18_669_WT_OF after

2

2

Example result folders. The output folder consists of one subfolder for each of the mice/signal that were included in the analysis, as well as a folder named “Combined information and comparison results” that contains combined signal figures and group comparison results.

### Result Demonstration
### Signal Processing

1

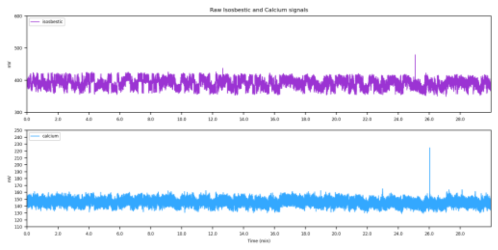


2

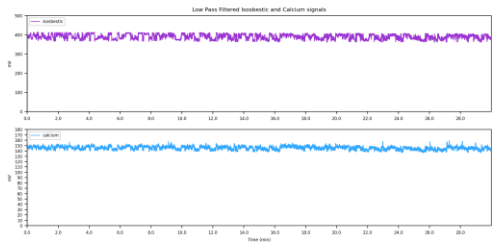


3

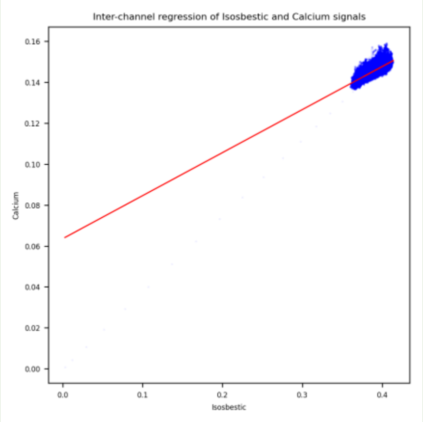


4

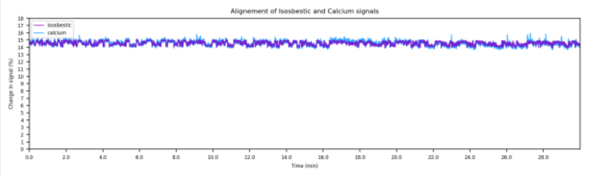


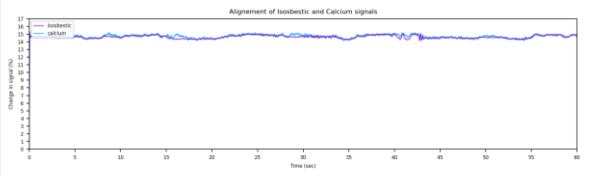


5

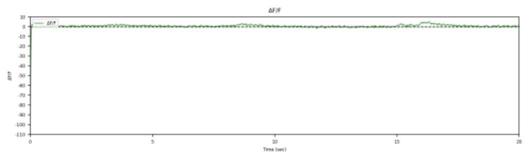


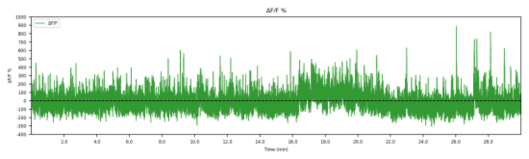


Figure results from individual signal file folders for the signal processing steps are demonstrated here.

1 Example of raw fiber photometry signals.

2 Example of fiber photometry signals after filtering separately.

3 Example of the isosbestic channel being fitted to the dopamine/calcium channel using least squares linear regression.

4 Example of overlapping the two channels after the fitting for 20s, 60s, and the whole during of the experiment.

5 Example of the calculated percent dF/F signal

### Result Demonstration
### Behavior Bout and Tracking Video

1

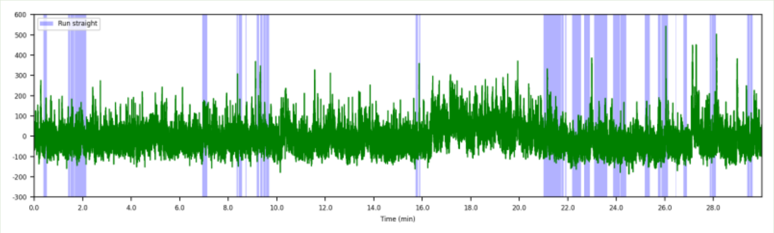


2

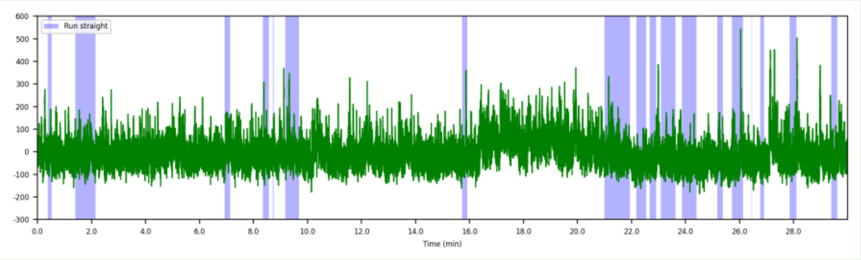


3

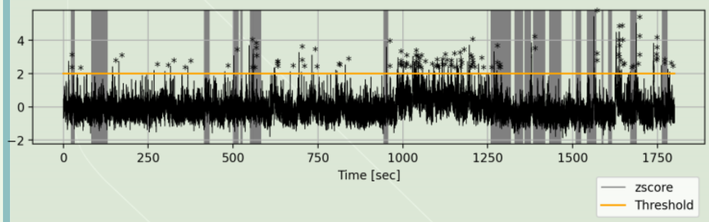


4

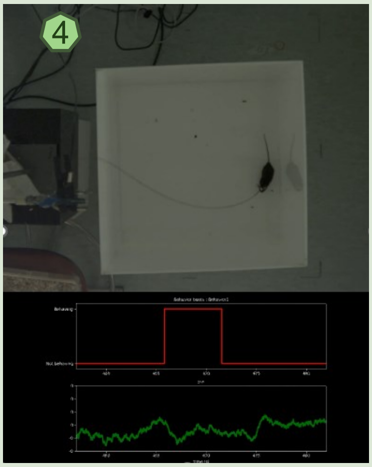

1 Example of the signals combined with the behaviour bouts, before making any user defined changes.

2 Example of the signals integrated with the behaviour bouts, after implementing the user defined minimum bout lengths and the minimum bout distances.

3 Example of the signals integrated with the behaviour bouts and the peak detection thresholding. The stars above the peaks show the peaks that are detected after thresholding.

4 Example of output video files. The video output files has the FP signals and the behaviour bouts integrated.

### Result Demonstration
### Comparison of Behavior Bouts

1

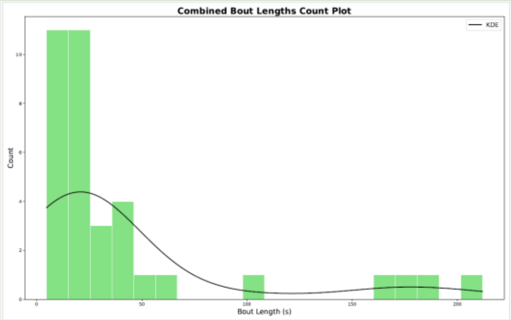


2

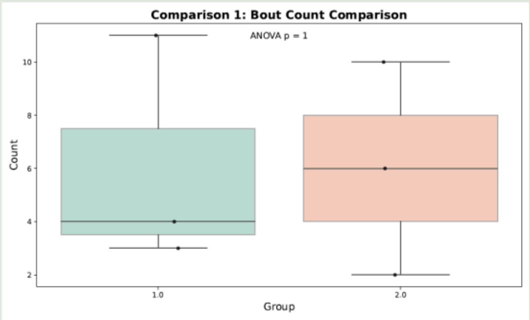


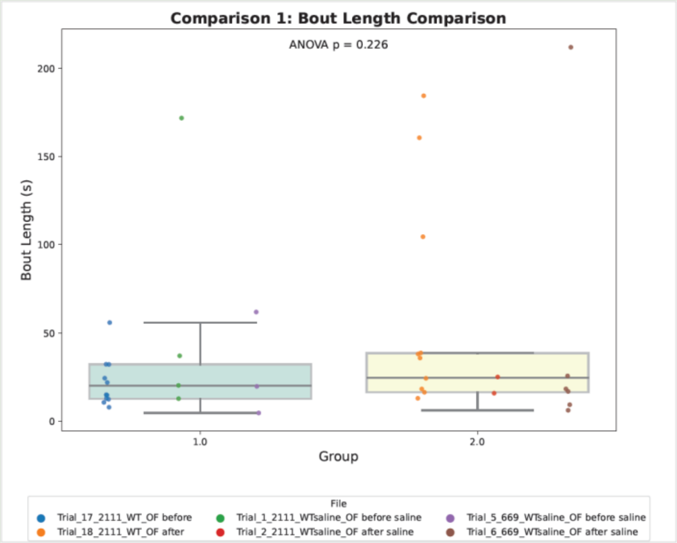


3

Results from this section focuses on bout analysis; this is the only result section which analyzes behaviors only without signals.

1 Here is an example of a histogram for behavior bout lengths across all files. A density estimate curve calculated with Gaussian kernel is overlaying the histogram. This plot is done for each behavior analyzed. Results (pdf and tiff formats) are named in the following format: combined_boutLength_histogram_for_[Walk straight].

2 Here is an example of a boxplot for bout count distribution of each comparison group. Each point is a bout count value from one experiment. Results (pdf and tiff formats) are named in the following format: comparison_[1]_bout_count_comparison_for_[Walk straight].

3 Here is an example of a boxplot for bout length distribution of each comparison group. Points are colored by their respective experiment. Each point represents the bout length of a single behavior bout, and all behavior bouts from each experiment are included. Results (pdf and tiff formats) are named in the following format: comparison_[1]_bout_length_comparison_for_[Walk straight].

*content in the square brackets are different for different comparisons and behaviors.

### Result Demonstration
### Comparison on Full Signal Level

1

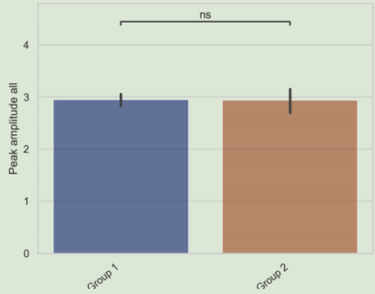


2

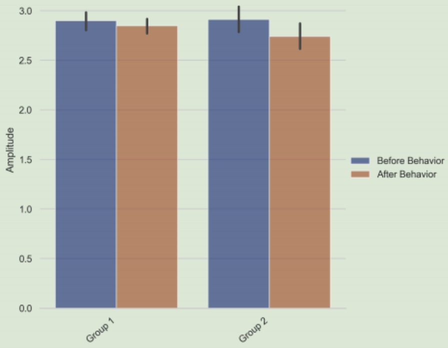


3

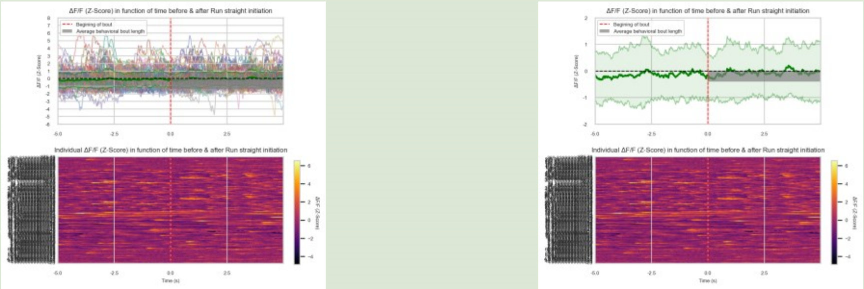

4

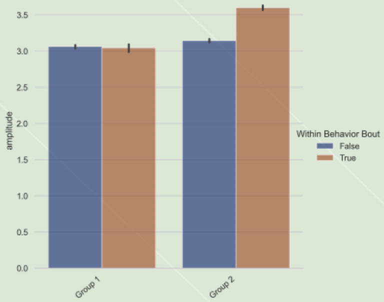


Signal peak data are analyzed on three levels: whole file average, block average, and individual peak level. For whole file averages, each peak property's average value per file is used in statistical tests and visualized in bar plots with T-test significance.

1 Here overall amplitude is shown and plotted between 2 groups. Results (png and svg formats) are named as “comparison1_Fullrange_Peak amplitude all_for_behaviour”

2 Here, the amplitude of the 5 seconds before and after the start of the behaving window is plotted as histograms and compared between 2 groups. The time plotted (5 sec in this example) is user defined. Results (png and svg formats) are named as “comparison1_behavior_range_Amplitude_for_behaviour”.

3 Heatmaps are plotted on the pooled behavior bout data. Results (in png and svg formats) are named as “comparison[1]_[compareGroup_1.0]_Peri_Event_Plot_for_[behaviour]_individualShown” and “comparison[1]_[compareGroup_2.0]_Peri_Event_Plot_for_[behaviour]_averageOnly”.

4 Here, the amplitude of the signals within and outside the behaviour windows are plotted. Results (in png and svg formats) are named as “comparison[1]_peak_behaviorVSnonbehavior_[amplitude]_for_[behaviour]”.

*content in the square brackets are different for different comparisons and behaviors.

### Result Demonstration
### Comparison on Time Block Level

1

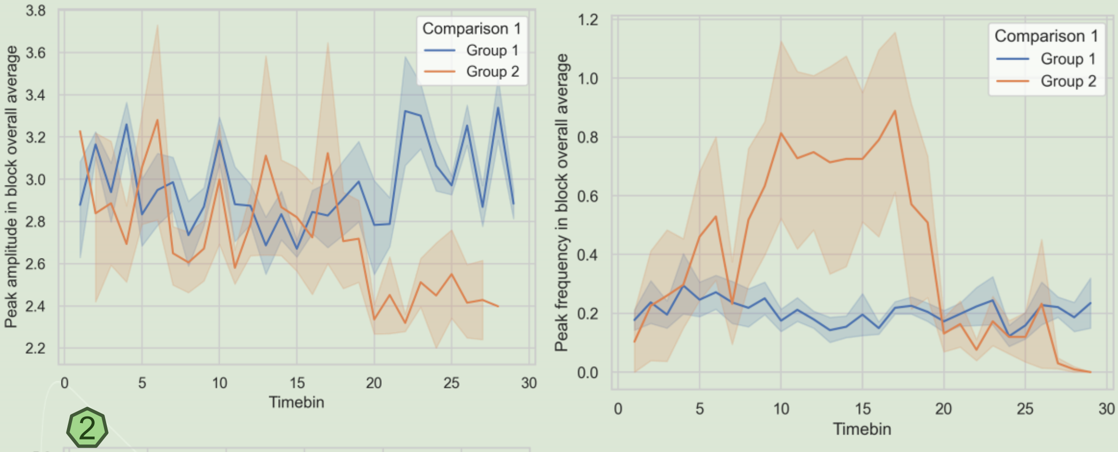


2

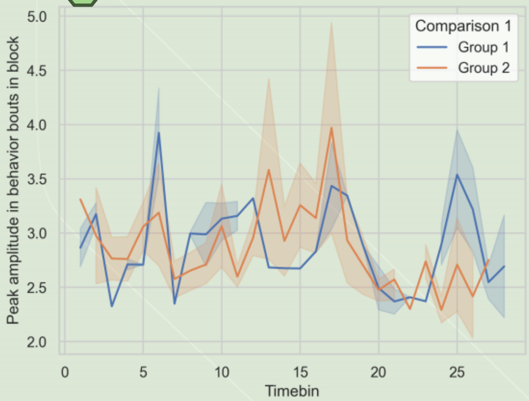


Signal peak data are analyzed on three levels: whole file average, block average, and individual peak level. For block averages, average values are calculated per block, and line plots display these averages by group.

1 Here the results (amplitude and frequency) are plotted on a block level. The time blocks here is 60 seconds (user defined) and the comparison is between 2 groups (based on input groups). The peaks are calculated from all the behaving and non behaving windows. Results (png and svg formats) are named as “comparison[1]_block_[Peak amplitude in block overall average]_for_[behaviour]”

2 Here the results (amplitude and frequency) are plotted on a block level. The time blocks here is 60 seconds (user defined) and comparison is between 2 groups (based on input groups). The peaks are calculated from all the behaving windows. Results (png and svg formats) are named as “comparison[1]_block_[Peak amplitude in behavior bouts in block]_for_[behaviour]”.

*content in the square brackets are different for different comparisons, behaviors and peak parameters.

### Result Demonstration
### Comparison on Peak Level

1

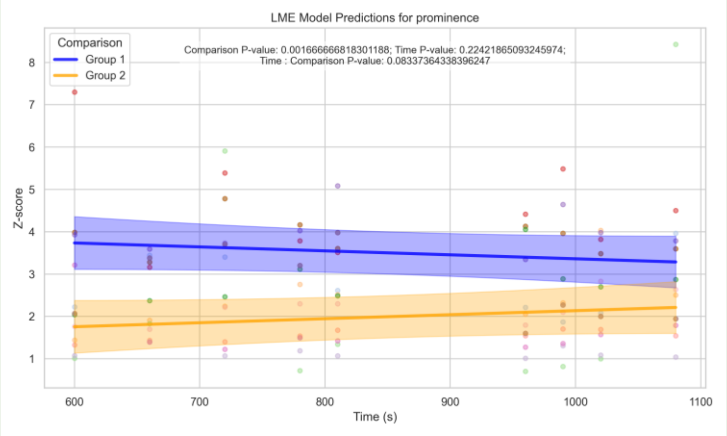


2

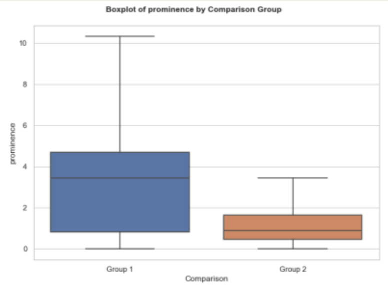


To account for repeated measures from the same subject in the same condition, the p-values are calculated from linear mixed effect models for each behavior analyzed. A scatterplot is used to display peak vs time relationship, and a boxplot is included for an overview of the peak distributions between comparison groups.

1 Here is an example of a scatterplot. Each point is a peak; the points are colored by experiment. The predicted linear models are overlaying the points, one per comparison group, and colored by comparison group. Each line has a confidence interval calculated based on standard error. The statistical information included came from LMEM. These results (pdf and tiff formats) are named in the following format:
Comparison[1]_singlePeaks_[amplitude]_lme_plot_[peaks_in_behaving]_[amplitude]_for_[Walk straight].

2 Here is an example of a boxplot. Each box is the distribution of the peaks from the respective compare group. Any statistical information included came from LMEM. These results (pdf and tiff formats) are named in the following format: comparison[1]_singlePeaks_[prominence]_boxplot_[all_peaks]_[prominence]_for_[Turn R]

*content in the square brackets are different for different comparisons, behaviors and peak parameters.

### Result Demonstration
### Segment Analysis – Before/After Bout

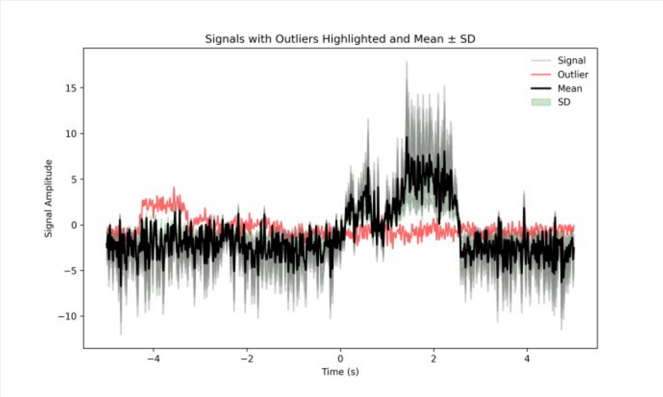


1

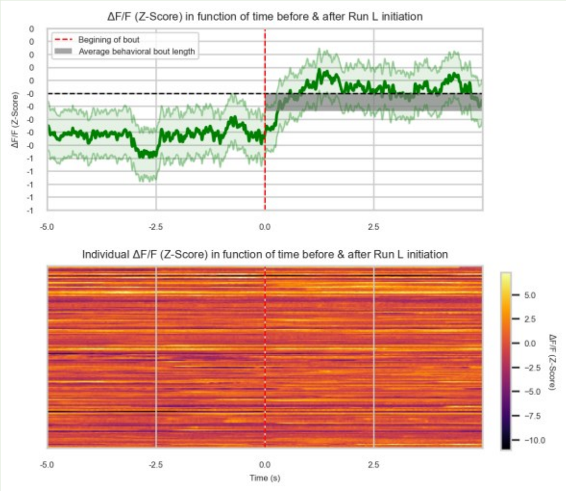


2

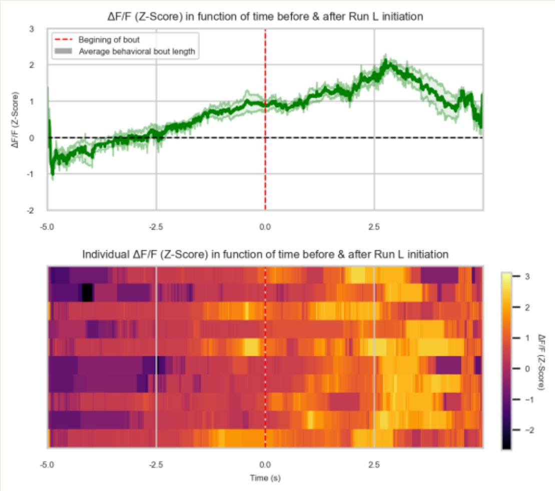


3

The segment analysis include several analysis steps, the first step is an optional outlier removal step, and the first figure shows the outliers detected in this step. A clustering step and an alignment step follow. For each cluster's alignment, several figures are plotted, the most representative peri plot is demonstrated here.

1 Here is an example of a line plot highlighting the outlier signals detected. Sometimes it's difficult to tell by eye the highlighted signals are outliers, as they may not have the extrema ranges of the signal distribution. The users should practice caution when determining whether the outlier removal step is suitable for their data. These results (tiff format) are named in the following format:
comparison[1]_compareGroup[1]_behavior_range_outlierID_for_[Walk straight]

2 3 Here are example peri plots. Figure 2 displays the signal before the alignment steps, and figure 3 displays the signal after the alignment steps. In Figure 2, the average signal is displayed with a standard error range. In figure 3, each cluster's aligned barycenter is displayed with a standard error range. Below is a heatmap displaying every individual signal. The time period for this analysis is determined by the user. The default setting is 5 seconds prior to the start of a behavior bout vs 5 seconds into the behavior bout. These results (png and svg formats) are named in the following formats: "comparison[1]_compareGroup_[1.0]_Peri_Event_Plot_for_[Running]_averageOnly" and "comparison[1]_compareGroup[2]_cluster[1]_afterAlign_Peri_Event_Plot_for_[Turn R]_averageOnly"

*content in the square brackets are different for different comparisons, comparison groups, and behaviors.

### Result Demonstration
### Before/After Bout Signal Analysis

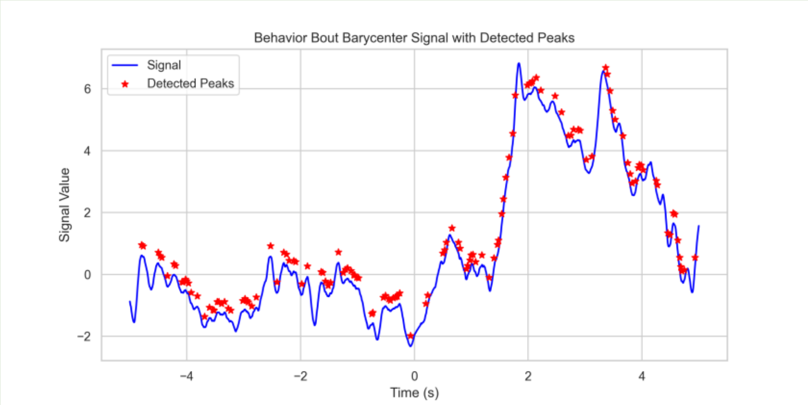


1

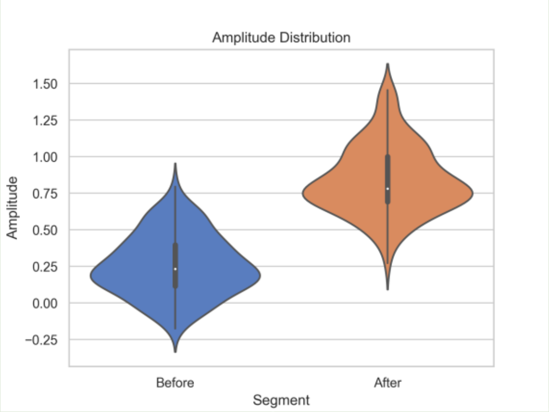


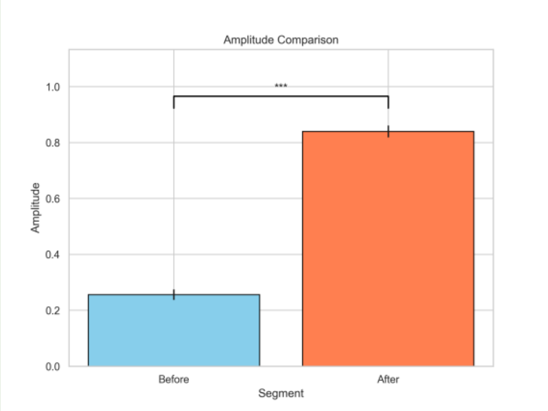


2

3

An optional follow up analysis can be performed for the aligned signal bouts. The barycenter for each behavior, comparison group, and cluster goes through peak extraction, and peak properties of before vs after behavior are compared.

1 Here is an example of extracted peaks on a barycenter of before/after bout signals. The * in red are the extracted peaks on the signal curve. The user can choose peak distance threshold for the peak extraction. These results (tiff and svg formats) are named in the following format:
comparison[1]_compareGroup[2]_cluster[1]_peakCompare_for_[Walk straight]_full_peaks

2 3 Here are the result figures for comparing before and after behavior bout signals. Signals are pooled into before behavior bout and after behavior bout distribution. Figure 2 shows a violin plot demonstrating the distributions, and figure 3 shows a boxplot with the same content. Wilcoxon test is used to calculate the p-value showing in the boxplot. All peak properties are analyzed. These results (tiff and svg formats) are named in the following formats:
“comparison[1]_compareGroup[2]_cluster[1]_peakCompare_for_[Walk straight]_[Prominence]_violinplot” and
“comparison[1]_compareGroup[2]_cluster[1]_peakCompare_for_[Walk straight]_[Frequency]_barplot”.

*content in the square brackets are different for different comparisons, comparison groups, clusters, behaviors, and peak properties.

### Result Demonstration
### Comparison for Two Conditions

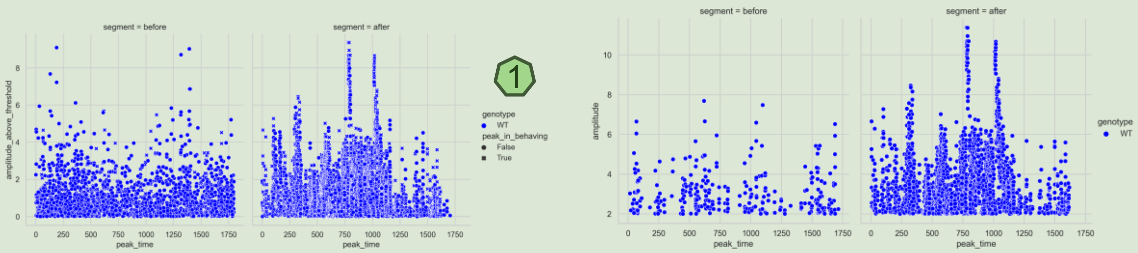


1

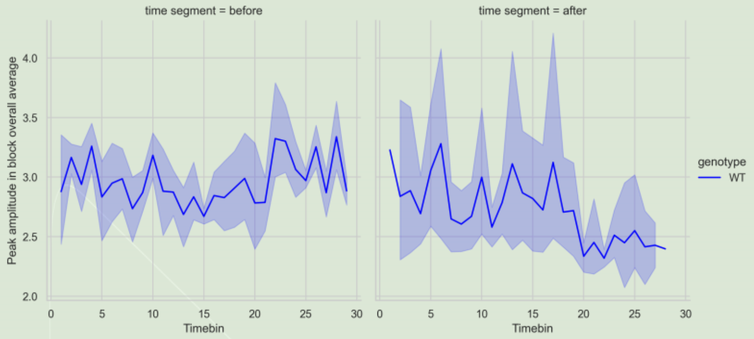


2

A default comparison setup allows comparing two conditions from the file information: one genotype and one condition (e.g., drug treatment). Users can switch the genotype information if needed. This analysis is optional.

1 2 Segment line plots and scatter plots (with both white and transparent backgrounds) are generated for time block and single peak data of peak properties, segmented by the first condition and grouped by genotype.
